# Mismatch between bird species sensitivity and the protection of intact habitats across the Americas

**DOI:** 10.1101/2021.03.28.437197

**Authors:** Victor Cazalis, Megan D. Barnes, Alison Johnston, James E.M. Watson, Cagan H. Şekercioğlu, Ana S.L. Rodrigues

**Affiliations:** CEFE, Univ Montpellier, CNRS, EPHE, IRD, Montpellier, France; German Centre for Integrative Biodiversity Research (iDiv) Halle-Jena-Leipzig, Puschstr. 4, D-04103 Leipzig, Germany; Leipzig University, Ritterstraße 26, 04109 Leipzig, Germany; Centre for Environmental Economics and Policy, School of Agriculture and Environment, University of Western Australia, Crawley, Western Australia, Australia; Cornell Lab of Ornithology, Cornell University, 159 Sapsucker Woods Road, Ithaca, NY, 14850, USA; Centre for Biodiversity and Conservation Science, School of Earth and Environmental Sciences, University of Queensland, St Lucia, Queensland 4072, Australia; Wildlife Conservation Society, Global Conservation Program, Bronx, New York 10460, USA; University of Utah, School of Biological Sciences, 257 South 1400 East, Salt Lake City, UT 84112, USA; Koç University, Department of Molecular Biology and Genetics, Rumelifeneri, Sarıyer, İstanbul, Turkey

## Abstract

Protected areas, the most prevalent international policy mechanism for biodiversity conservation, are highly heterogeneous in their effectiveness at buffering ecosystems and species’ habitats from human pressure. Protected areas with intense human pressure cannot protect species that are highly sensitive to human activities. Here, we use 60 million bird observations from the eBird citizen science platform to estimate the sensitivity to human pressure of each bird species breeding in the Americas (Nearctic and Neotropical regions). We find that high-sensitivity species, while found in all ecoregions, are concentrated in the tropical biomes. Ecoregions with large proportions of high-sensitivity species do not have more intact protected habitat, resulting in a low coverage of intact protected habitat for many high-sensitivity species. What is more, 139 high-sensitivity species have little or no intact protected habitat within their distributions while being threatened with extinction. Finally, we show that protected area intactness is decreasing faster in ecoregions with many high-sensitivity species. Our results highlight a major mismatch between species conservation needs and the coverage of intact protected habitats, and will likely hamper the long-term effectiveness of protected areas at retaining species. We highlight ecoregions where the protection and management of intact habitats, complemented by the restoration of degraded ones, is urgently needed to avoid extinctions.

## Introduction

Protected areas are clearly defined geographical spaces, recognised, dedicated and managed to achieve the long-term conservation of nature (UNEP-WCMC, IUCN and NGS, 2020). Acknowledged as one of the world’s most important biodiversity conservation tools (Watson et al., 2014; Maxwell et al., 2020), there has been a marked expansion in their extent over the past few decades and their known coverage currently reaches 15.4% of the global land surface (UNEP-WCMC, IUCN and NGS, 2021). Ongoing renegotiation of global policy targets are expected to result in a more ambitious 30% target (SCBD, 2020), likely to drive further expansion.

However, protected areas vary substantially in both their intended management (as legally defined) and in the practical effectiveness of their implementation (Geldmann et al., 2018; Barnes et al., 2016). As a result, many protected areas are currently under intense and increasing human pressure (Jones et al., 2018; Venter et al., 2016a). For this reason, the percentage of area protected is on its own insufficient to evaluate protected area effectiveness (Rodrigues and Cazalis, 2020). A pure focus on protected area expansion, without guarantee of concomitant quality, has been criticised as encouraging fast expansion into areas of little value to biodiversity, or with little on-the-ground implementation effort (Barnes et al., 2018; Visconti et al., 2019).

This high level of degradation of some protected habitats may be a concern in some cases because degradation is known to impact many species (Di Marco et al., 2019; Watson et al., 2018; Gibson et al., 2011). However, it does not necessarily follow that all protected areas need to be in perfectly intact condition. Indeed, while many species are highly sensitive to human pressure (Barlow et al., 2016; Gibson et al., 2011), many others can tolerate some levels of, or even benefit from, anthropogenic land use change (Guetté et al., 2017; McKinney, 2006; Şekercioğlu et al., 2019). Given the need to reconcile biodiversity conservation and human development, two broad strategies have been proposed: land sparing, focused on setting aside intact habitats while concentrating human pressure elsewhere; and land sharing, integrating conservation and development by spreading human pressure across larger areas, including multiple use protected areas (Green et al., 2005; Phalan et al., 2011; Williams et al., 2017). In practice, the best strategy for each species depends on how it responds to human pressure (Green et al., 2005). Species that respond negatively to even low levels of pressure require strict management of sufficient expanses of intact habitats (with a concomitant concentration of human activities elsewhere; i.e., land sparing), whereas species that can tolerate moderate to high human pressure may be adequately protected in multi-use protected areas, or not need protected areas at all (i.e., land sharing). The few studies quantifying species responses to increasing human pressure (based on gradients of agriculture yield or urban intensification) in the context of the land sparing/land sharing debate found that many species are strongly impacted by even low levels of human pressure, providing support for the need to set aside intact areas for their conservation (Phalan et al., 2011; Williams et al., 2017; Collas et al., 2017; Şekercioğlu et al., 2019). Understanding how these sensitive species are distributed across the world may in turn shed light on the extent to which different regions are more or less dependent on the protection of intact habitat for the conservation of their biodiversity.

Here, we investigate the large-scale spatial variation in sensitivity to human pressure for 4,424 bird species, contrasting this with the distribution of intact protected habitat to highlight priorities for expanding or reinforcing intact protected habitat coverage. Taking advantage of the millions of field records collated through the eBird citizen science platform, we focus on the Americas as a study region. We first measure the sensitivity of 2,550 breeding species by modelling the relationship between abundance and human pressure as measured by the human footprint index. We then impute the sensitivity of the remaining 1,874 breeding species based on their traits. Using ecoregions as spatial unit, we contrast spatial patterns in species sensitivity with the coverage of intact protected habitat of ecoregions, in order to identify the regions with a critical mismatch between the two. In addition, we identify as species of major concern those that are simultaneously highly sensitive to human pressure, whose distributions have minor or no coverage by intact protected habitats, and that are globally threatened with extinction. Finally, we analyse these results in light of the recent trends in human footprint to understand if the adequacy of protected areas to the conservation of high-sensitivity species is improving or worsening.

## Methods

### Study area

We focused on the Americas (i.e., Nearctic and Neotropical realms, covering the North and South American continents) given the large concentration of bird observations and wide ecological variation (major latitudinal gradients, representing all major biomes). We analysed data within the corresponding 325 terrestrial ecoregions (Olson et al., 2001) for which there is human footprint data (Figs. 2-3).

### Species data

#### Bird distribution and abundance data

We used data from eBird (Sullivan et al., 2014), a unique online platform gathering hundreds of millions of bird observations across the globe, with a semi-structured data collection framework. Observers report their observations in checklists and provide information on sampling effort, allowing eBird data to be transformed into a dataset enabling estimates of indices of relative abundance (Sullivan et al., 2009), as described below. We used the eBird dataset released in October 2019 (eBird, 2019).

In addition, we used species distribution data (coarse polygons delimiting the species’ ranges) as mapped by BirdLife International and HBW (2019), having aligned the taxonomies between these two datasets by following the latter (Supplementary Methods 1).

#### Building a standard abundance dataset

We filtered the dataset following guidelines provided by the eBird team (Sullivan et al., 2009; Strimas-Mackey et al., 2020; Johnston et al., In press), similarly to the data filtering process described in Cazalis et al., (2020). We restricted our dataset to recent (2010-2019) observations, to increase synchronisation between bird records and landscape data. We kept only checklists for which observers certified having reported all species identified, thus obtaining a count dataset that included non-detections. Further, we filtered checklists based on sampling effort and protocol to create a set of checklists with relatively consistent effort. Specifically, we kept only checklists that reported a duration of sampling between 0.5-10 hours and less than 5 km distance travelled arising from: the ‘Stationary Points’ protocol (i.e., the observer did not move during sampling); the ‘Travelling’ protocol (i.e., the observer moved during sampling); or ‘Historical counts’ (if they included information on duration and distance travelled). This protocol selection excludes sampling events targeting particular taxonomic groups (e.g. wader surveys, nocturnal protocols) or using specific methods (e.g. banding). Each checklist is associated with a point, with coordinates provided by observers (often the middle of the route for ‘Traveling’ checklists).

Pseudo-replication in the database may occur when several observers record birds at the same place at the same time. To eliminate these records, we first used the *auk_unique* function from the *’auk’* package (Strimas-Mackey et al., 2017), specifically designed to process eBird data. This function combines checklists from multiple observers reported to be birding together, in order to obtain a single checklist for each sampling event (the number of observers associated with each checklist alters the detection probability, and will be subsequently controlled for). Second, because even observers who are not birding together can create pseudo-replication if they overlap in space and time, whenever several checklists were less than 5 km apart on the same day we randomly and sequentially selected one checklist (i.e., iteratively until no two checklists were less than 5 km apart). All the checklists we selected were ‘complete’ with each observer recording all the species they could detect and identify. However, more experienced observers will be able to detect and identify more species and therefore non-detections will be more likely to equate to real species absences. Therefore, we only considered observations made by observers with a minimum level of experience, i.e. those who submitted ≥ 50 checklists, including ≥ 100 species during the study period (2010-2019; see Supplementary Methods 2).

We excluded from the dataset “not approved” observations (corresponding to exotic, escaped or feral individuals), as well as domestic species, but kept established invasive species. Using the *auk_rollup* function from the *’auk’* package, we removed subspecies, moving all observations to the species level. We also excluded marine species, defined as species with ‘sea’ or ‘coastal’ as primary habitat (cf. bird species traits section below).

In many species, individuals are the most territorial and selective in terms of habitat requirements when breeding (Zuckerberg et al., 2016). We thus focused our analyses on potentially breeding individuals, by narrowing each species’ observations records to the respective breeding season and breeding grounds. For each 10° latitudinal band, we derived the broad breeding season (all species considered together) based on the temporal distribution of records coded as ‘breeding’ in the eBird database (e.g., 6 April to 9 August for latitudes 50°N to 60°N; all year round for latitudes 10°S to 10°N; see Supplementary Methods 3). Within these dates, we then focused on the observations made in the breeding grounds of each species (as mapped by BirdLife International and HBW (2019)).

We considered a species absent if undetected in checklists made during the breeding season and located within the breeding range of the species. For species recorded as present, we used the count (i.e., number of individuals observed) provided by observers. In some cases (4% of the observations in our dataset) observers did not provide an abundance and instead reported species presence with an “X”. We treated these as *NA* values in our analyses.

The final dataset in our main analyses consisted of 59,583,879 observations and 404,086,397 inferred absences, structured into 3,449,486 checklists made by 44,013 observers and representing 4,424 species.

#### Accounting for observer differences

Even when the sampling protocol is similar, eBird checklists may greatly differ because of the important heterogeneity in observers’ experience, skills, behaviour, and equipment. As mentioned above, observations were filtered to include only those made by observers with a minimum level of experience. In addition, we calculated an individual observer calibration index, which we included as control variable in all subsequent analyses of eBird data. This index (closely related to the one calculated in Cazalis et al., (2020), following Kelling et al., (2015) and Johnston et al., (2018)), uses a mixed model with random effect of observer to estimate the log-scaled number of species each observer is expected to report in an average sampling event (see details in Supplementary Methods 2).

#### Bird species traits

For each of 4,424 species considered, we obtained data on eight species-level variables (that we call “traits” for simplicity): four categorical (primary habitat, primary diet, migratory status, and taxonomic Order), and four continuous (body mass, specialisation, Red List status, and breeding range size in the Americas). Trait data came from two sources: the trait database BirdBase, and BirdLife International datasets.

BirdBase is a regularly updated global database of the ecology and life history traits of the world’s bird species (Şekercioğlu et al., 2004, 2019). We extracted from it the species’ primary habitat, structured into 10 classes (after exclusion of ‘coastal’ and ‘sea’ species): Artificial, Deserts, Forests, Grasslands, Riparian, Rocky, Savannahs, Shrublands, Wetlands, Woodlands. Primary diet consisted of eight classes (after combining ‘Carnivore’, ‘Scavenger’ and ‘Vertebrate’ into ‘Carnivorous’; combining ‘Plant’ and ‘Herbivore’ into ‘Herbivorous’; and considering the 42 species with an ‘unknown’ diet as ‘Omnivorous’): Carnivorous, Frugivorous, Granivorous, Herbivorous, Insectivorous, Nectarivorous, Omnivorous, Piscivorous. In addition, we obtained from the same database: migratory status (‘strict’, ‘partial’, or ‘sedentary’); body mass; and species taxonomic Order (e.g., Accipitriformes, Anseriformes). We calculated a specialisation index for each species based on the number of different habitat (HB) and diet (DB) categories for each species, as in Şekercioğlu (2011): ln [100/(HB × DB)]. We inferred specialisation and body mass values of species for which it was unknown (respectively 43 and 358), by using the mean specialisation and mass of the documented species in the same taxonomic Family.

We extracted Red List status from BirdLife International (2019), and (following Butchart et al., 2007) transformed it into a quantitative variable: Least Concern (LC) as 1, Near Threatened (NT) as 2, Vulnerable (VU) as 3, Endangered (EN) as 4, and Critically Endangered (CR) as 5. We treated the six species for which the Red List status was Data Deficient as LC. We calculated each species’ breeding range size (as defined above) from BirdLife International and HBW (2019), within our study area.

### Landscape data

#### Protected areas

We used spatial protected area data from the World Database on Protected Areas (UNEP-WCMC & IUCN, 2020), following the standard filtering procedure (UNEP-WCMC and IUCN, 2019) that excludes ‘Man and Biosphere’ reserves, protected areas with no associated polygons and those that are not yet implemented (i.e., we kept only those ‘designated’, ‘inscribed’, or ‘established’).

#### Human footprint

We used maps of human footprint in 2000 and in 2013, in raster format at a resolution of ∼1km (Williams et al., 2020). These maps are an updated and more complete version of the index derived by Venter et al. (2016b), generated from the combination of eight human pressure variables (built environments, population density, night-time lights, crop lands, pasture lands, accessibility via roads, railways and navigable waterways) and ranging from 0 (perfect intactness) to 50 (extremely high pressure). Human footprint data have previously been used to analyse species’ responses to human pressures (Di Marco et al., 2018; Barnagaud et al., 2019; Cazalis et al., 2020).

We assigned to each checklist a value of human footprint in 2013 by calculating the mean value of human footprint in pixels intersecting by at least 1% a buffer around the checklist coordinates. We considered a buffer of 2.5 km radius to ensure it covers the large majority of the area sampled by the selected travelling protocols.

#### Altitude

The altitude of each checklist was calculated as the mean altitude of all pixels from the Global Land One-kilometer Base Elevation raster (National Geophysical Data Center, 1999) intersecting by at least 1% the 2.5 km buffer around the checklist coordinates.

#### Net Primary Productivity

We calculated a Net Primary Productivity (NPP) value for each checklist, using NASA’s Earth Observatory Team (2020) maps. We first created a raster layer by calculating for each cell the mean of NPP values from January 2014 to November 2016 (December 2016 data were not available). We then extracted for each checklist, the mean value of each pixel in this raster that intersected by at least 1% the 2.5 km buffer around the checklist coordinates.

## Analyses

We analysed the data to answer four questions: (1) How sensitive are bird species to Earth’s terrestrial human footprint? (2) Where across the study area are species the most sensitive? (3) Is current protection of intact habitats matched to the species’ needs? (4) Are protected areas retaining intact habitats over time?

### How sensitive are species to human footprint?

#### Direct measure of sensitivity (data-rich species)

We quantified directly the sensitivity to human footprint for a subset of 2,550 data-rich species, selected according to three conditions: ≥ 200 records with abundance; ≥80% of all records with abundance (as ‘X’ often concerns observations with too many individuals to be counted, which could introduce a bias for gregarious species); and with distributions across a wide range of human footprint values. The latter were selected by first calculating the 1% and the 99% quantiles of human footprint from all checklists within each species’ breeding distribution, and then keeping only those species for which these quantiles differed by ≥ 25.

For each of these 2,550 data-rich species, we ran a General Additive Model (GAM) modelling the link between species’ abundance and human footprint of the checklists’ location (assuming a negative binomial distribution of abundance and using the bam function; (Wood, 2011)). In order to enable a diversity of links, from linear to non-monotonous relations, we used a smoothed term on human footprint, but we constrained the degree of the smoothing function to 6 to avoid very complex functions (see examples of relations in Fig. 1). In these models, we controlled for differences in sampling effort (logarithm of sampling duration; logarithm of number of observers; observer calibration index, using the maximum if multiple observers), differences in ecological conditions (altitude and net primary productivity, both assuming parabolic responses; we could not use smoothed-terms here because of computing limitations), and large-scale patterns of spatial trends (interacting smooth-term with longitude and latitude), with the following structure:

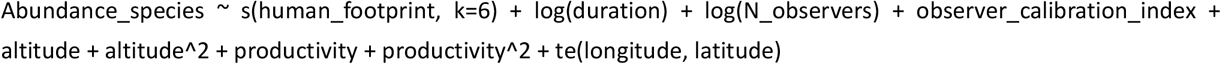

**Fig. 1:**
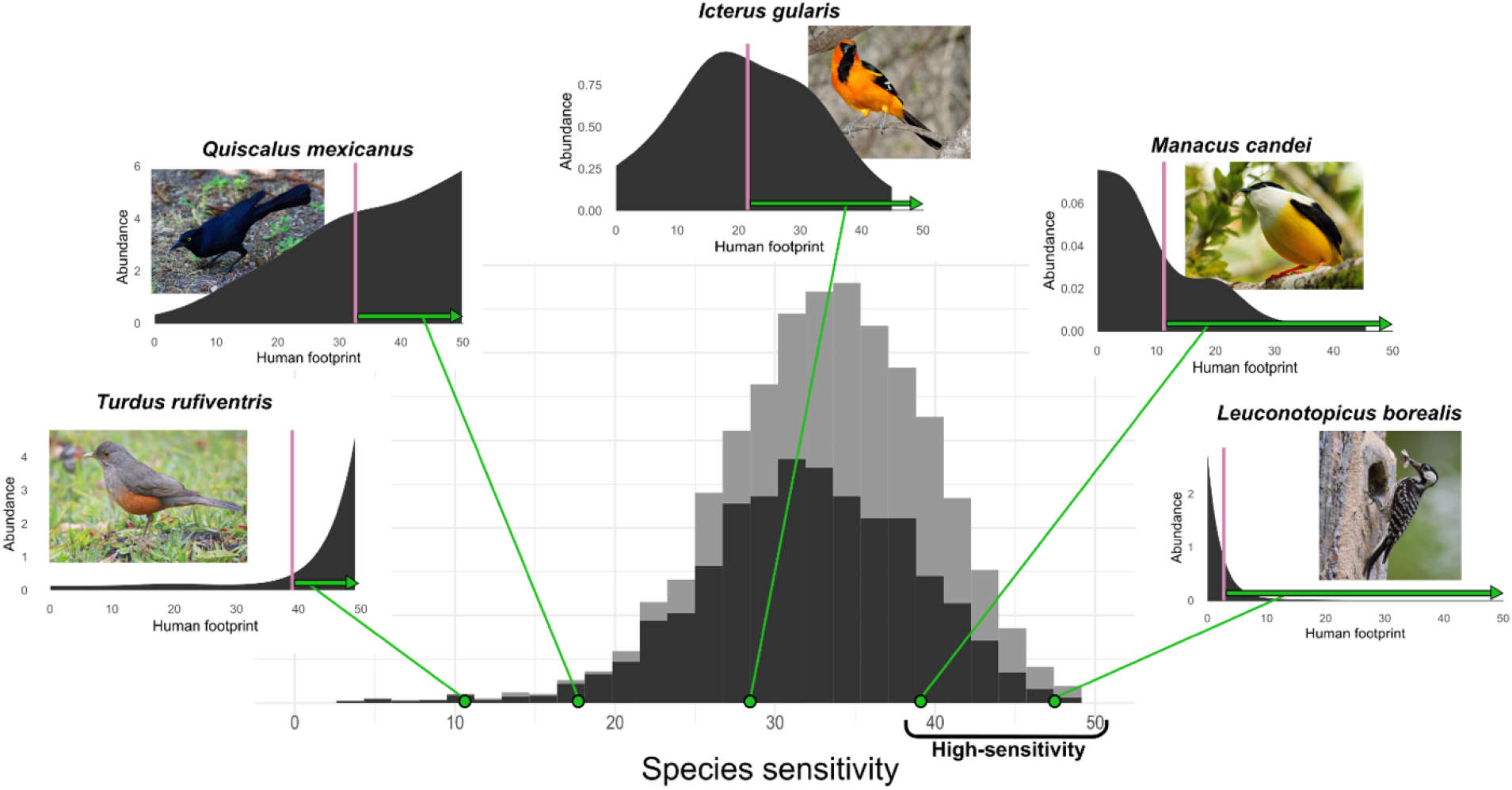
Stacked frequency distribution of sensitivity values across all 4,424 bird species that breed in the Americas, including data-rich species for which sensitivity was measured directly (dark grey, N=2,550), and data-poor species for which it was imputed from trait information (light grey, N=1,874). Insets correspond to five examples of data-rich species across a gradient of sensitivity, showing for each: the modelled response of abundance to human footprint; the measure of sensitivity (green horizontal arrow) obtained from the difference between 50 (i.e., the maximum value of human footprint) and the weighted mean value of predicted abundance (pink vertical line). High-sensitivity species were defined as the 25% most sensitive ones. Photo credits: *T*.*r*. (Luiz Carlos Rocha, https://www.flickr.com/photos/luizmrocha/), *Q*.*m*. (BarbeeAnne, https://pixabay.com/photos/cuba-black-bird-great-tailed-grackle-2555949/),*I*.*g*. (Skeeze, https://www.needpix.com/photo/download/729995/altamira-oriole-bird-perched-nature-wildlife-songbird-branch-outdoors-birdwatching), *M*.*c*. (David Rodriguez Arias,https://www.flickr.com/photos/82969027@N04), *L*.*b*. (Sam D. Hamilton, https://upload.wikimedia.org/wikipedia/commons/6/61/Picoides_borealis_-Mississippi%2C_USA_-feeding-8.jpg).

We then predicted the relative abundance of the species across a gradient of human footprint ranging from 0 to the maximum value of human footprint observed within the species’ distribution, with a step of 0.05 (fixing all other variables to their median values) and extracted the weighted average of this distribution (Fig. 1). Finally, we measured each species’ sensitivity as the difference between 50 (i.e., maximum human footprint) and this weighted average. High-sensitivity species have an abundance strongly biased towards sites with low human footprint (e.g., *Leuconotopicus borealis*), medium-sensitivity species have an abundance unrelated to human footprint or biased towards medium human footprint (e.g., *Icterus gularis*), and low-sensitivity species have an abundance biased towards sites with high human footprint (e.g., *Turdus rufiventris*) (Fig. 1).

#### Imputed sensitivity (data-poor species)

We used information on species’ traits to impute the sensitivity of the remaining (data-poor) 1,874 species, from the 2,550 data-rich species. To do so, we first linked the values of sensitivity measured for data-rich species to their traits using a linear model. We included in the model the species’ primary habitat, primary diet, specialisation (log-scaled), body mass (log-scaled), Red List status, range size (log-scaled), migratory status, and taxonomic Order, with the following structure:

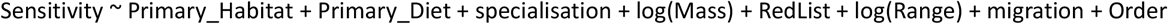

We then used this model to impute the sensitivity of the data-poor species (see details in Supplementary Methods 4).

We use species’ sensitivity as a relative measure, to compare among species and regions

(Supplementary Methods 5). We defined high-sensitivity species as the 25% most sensitive species (Fig. 1).

#### Where are bird assemblages the most sensitive?

As a measure of the sensitivity of bird assemblages to human footprint, we calculated for each ecoregion the proportion of all breeding species classified as high-sensitivity. The list of breeding species per ecoregion was obtained by overlapping species’ breeding distributions with the ecoregion’s boundaries.

We analysed whether our results were driven by methodological choices through two additional analyses. In the first one, we considered only data-rich species (N=2,550), of which 25% (N=638) were classed as high-sensitivity, and mapped the proportion of high-sensitivity species per ecoregion. In the second, again using all species (N=4,424 species), we mapped two alternative measures of the sensitivity of bird assemblages to human footprint: median sensitivity across all species that breed in the ecoregion; and absolute number of high-sensitivity species per ecoregion.

#### Is intact habitat protection matched to species’ needs?

We created a 0/1 raster layer of intact protected habitat across the study area as a transformation of the 2013 human footprint raster, by assigning the value 1 only to those pixels with intact habitat (human footprint value <4, Williams et al., 2020) whose centre was located within a protected area. We then used this layer to investigate if intact habitat protection is matched to species’ conservation needs (i.e., their sensitivity), both across space (ecoregions) and across species.

#### Across ecoregions

For each ecoregion, we measured the level of intact habitat protection as the proportion of its pixels (those included by >50% within the ecoregion) classified as intact protected habitat. We then analysed how this index relates to the proportion of high-sensitivity species per ecoregion, both quantitatively (using the Pearson correlation coefficient) and visually (mapping the correspondence between these variables using a bivariate colour scale).

For comparison, we also analysed two alternative measures of the investment in habitat protection per ecoregion: protected area extent (the fraction of the ecoregion covered by protected areas); and protected area intactness (the mean intactness of protected pixels, where intactness is the opposite of human footprint).

#### Across species

For each high-sensitivity species, we quantified their coverage by intact protected habitat. Given that a pixel measures ∼ 1 km^2^, we calculated the area of intact protected habitat per species by summing the values in the 0/1 raster layer of intact protected habitat for pixels included (by >50%) within the species’ breeding distribution.

We then assessed whether high-sensitivity species were adequately covered by intact protected habitat, by comparing their coverage by intact protected habitat against a predefined representation target. For each species, this target was calculated based on a widely-used approach (e.g., Rodrigues et al., (2004), Maxwell et al., (2020), Butchart et al., (2015)) whereby species with very small ranges (< 1000 km^2^) have a 100% target, those with very widespread ranges (>250,000 km^2^) have a 10% target, with the target for species with ranges of intermediate size being interpolated between these two extremes. A high-sensitivity species was considered inadequately covered if its coverage by intact protected habitat falls below this representation target. Among these, we highlight a subset we designate as species of major concern, defined by having no or minor coverage (≤ 10% of their target met; i.e., less than 10% of the range for highly-restricted species, less than 1% for species with very large range), as well as being globally threatened with extinction (Vulnerable, Endangered, or Critically Endangered).

### Are protected areas retaining intact habitats over time?

We created a raster layer that maps the trends in human footprint between 2000 and 2013, from the difference between human footprint values in 2013 and in 2000. We then calculated, for each ecoregion, the trend in protected area intactness as the mean decrease in human footprint in all pixels intersecting (by 50%) protected areas (i.e., a positive value means an increase in protected area intactness). Finally, we calculated, across ecoregions, the Pearson correlation coefficient between trends in protected area intactness and the proportion of high-sensitivity species.

## Results

### How sensitive are species to human footprint?

Across the 4,424 bird species that breed in the Americas, values of sensitivity to human footprint range from 2.9 to 50.0, following a Gaussian distribution with a median value of 33 (Fig. 1). Sensitivity is higher in data-poor species than in data-rich ones. The threshold for high-sensitivity species is 37.6. Among the 1,106 high-sensitivity species, 24.0% are threatened with extinction (compared with 7.0% for species not categorised as high-sensitivity).

### Where are bird assemblages the most sensitive?

Assemblages most sensitive to human footprint (i.e., with highest proportion of local species being high-sensitivity) are concentrated in tropical ecoregions, especially along the Andean mountain range and its eastern slopes towards the Amazonian basin, as well as in Central America (Fig. 2A).

**Fig. 2:**
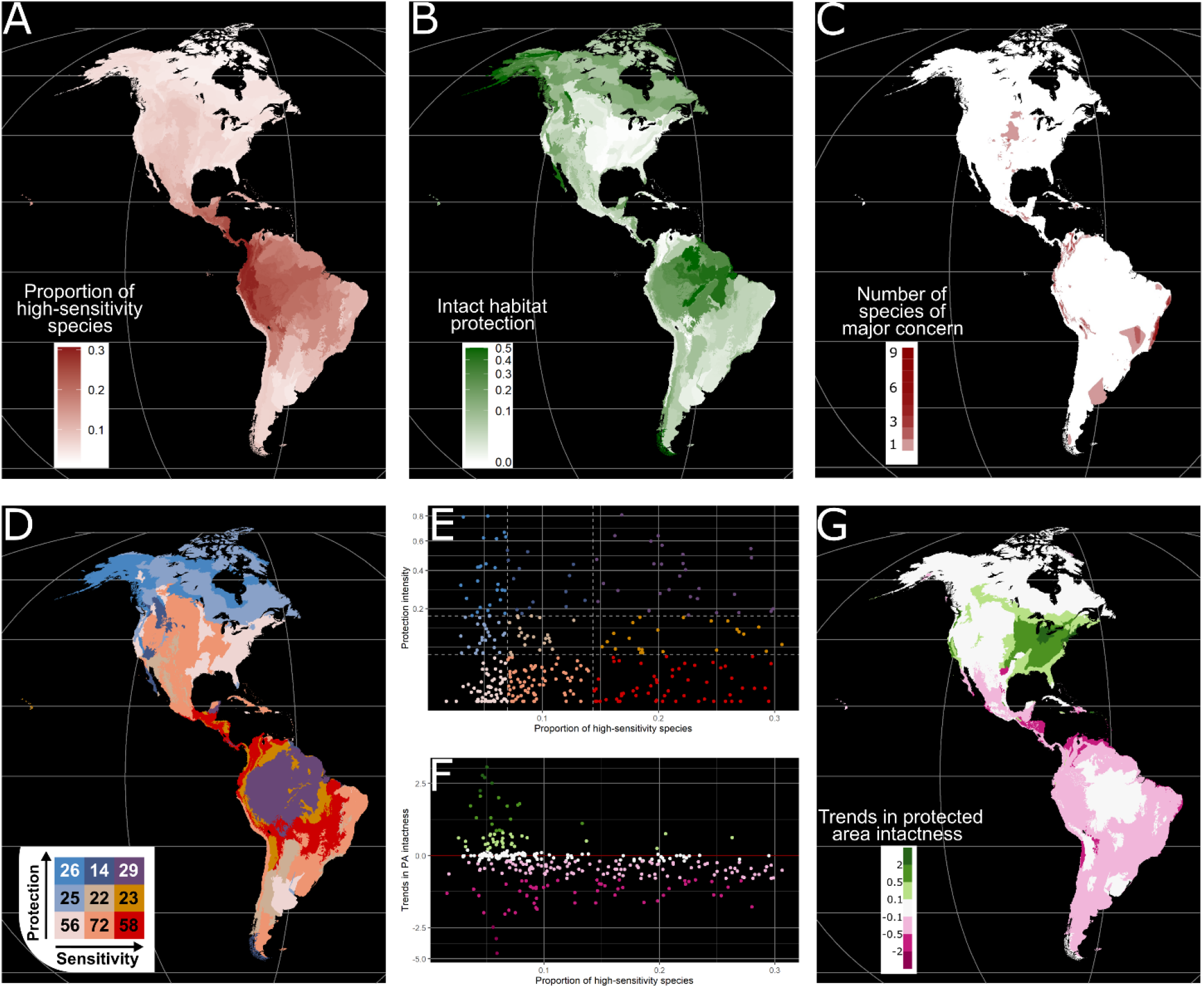
Spatial patterns in bird assemblage sensitivity and intact habitat protection. (A) Assemblage sensitivity measured as the proportion of species per ecoregion that are high-sensitivity species. (B) Intact habitat protection per ecoregion (i.e., proportion of ecoregion area simultaneously protected and with human footprint <4). (C) Spatial distribution of the 139 species of major concern, i.e., simultaneously high-sensitivity, threatened and with no or minor coverage by intact protected habitat. (D) Spatial pattern and (E) scatterplot of the relationship between assemblage sensitivity and intact habitat protection, across ecoregions. The bivariate colour scale in D and E is built by cutting proportion of high-sensitivity into terciles ([0; 0.070[,[0.070; 0.143[,[0.143; 1]) and intact habitat protection into [0; 0.05[, [0.05; 0.17[, [0.17; 1] (dashed lines in E). (G) Trends in protected area intactness per ecoregion (i.e., decrease in human footprint within protected areas between 2000 and 2013); and (F) in relation to the proportion of high-sensitivity species per ecoregion. Green shades show improvement, rose shades degradation.

Very similar patterns were found when considering only data-rich species (coef = 0.97, P < 2×10^−16^; Fig. S6), or when assemblage sensitivity was measured as the number of high-sensitivity species or the median sensitivity of species per ecoregion (Fig. S5). We thus focus henceforth on the proportion of high-sensitivity species measured across all species (Fig. 2A).

### Is intact habitat protection matched to species sensitivity?

#### Across ecoregions

Intact habitat protection is highest in the Amazonian basin, Boreal region, Western North America, and Patagonia (Fig. 2B, Table S2), where ecoregions combine relatively high protected area extent and high protected area intactness (Fig. S7). Conversely, intact habitat protection is lowest in Eastern

North America, and in much of South America outside the Amazonian basin (Fig. 2B). The proportion of high-sensitivity species per ecoregion is not correlated with its levels of intact habitat protection (coef = 0.044, P=0.424; Fig. 2D-E), neither with protected area extent, nor with protected area intactness (Fig. S7), and was not impacted by our measure of assemblage sensitivity (Fig. S5).

Areas with highly sensitive bird assemblages but low intact habitat protection are concentrated in tropical ecoregions, especially in the tropical Andes and their western slopes towards the Pacific coast, Colombia’s Choco region and Magdalena Valley, Venezuela Coastal Range, Central America, and the Cerrado savannahs of Brazil (red in Fig. 2D, Fig. 3). These ecoregions mainly correspond to tropical and subtropical forest biomes, mostly moist broadleaf (22 ecoregions), but also dry broadleaf (15), mangroves (7) and coniferous forests (3). They also include grasslands and shrublands, including montane (6), deserts and xeric (3), tropical (1), and flooded (1) (Fig. 3; Olson et al. 2001).

**Fig. 3:**
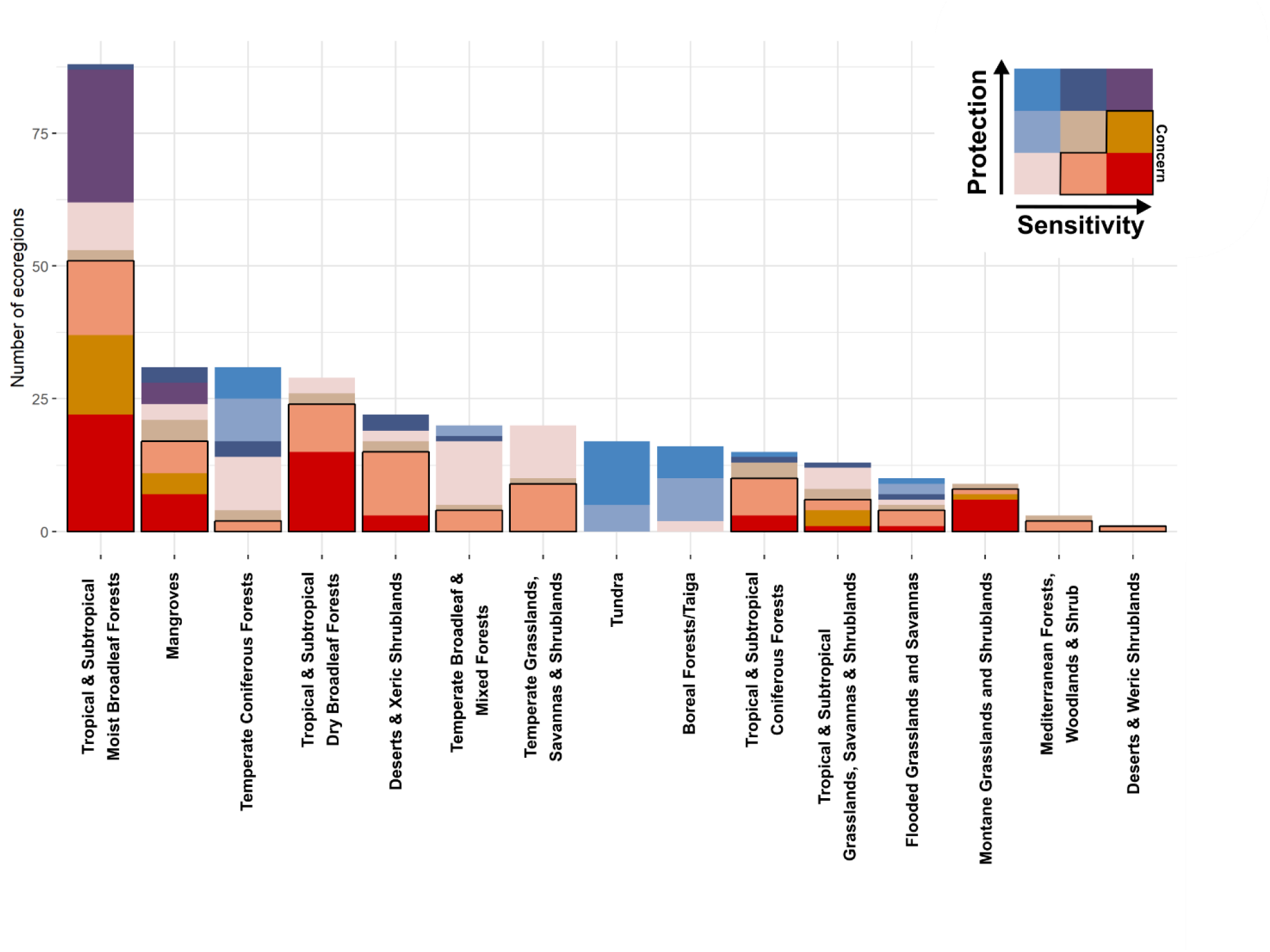
Distribution of ecoregions per biome (following Olson et al. 2001), colour-coded according to the relationship between the proportion of high-sensitivity species and intact habitat protection (colours as in Fig. 2D-E). The three categories of particular concern are outlined in black.

Additionally, 23 ecoregions have high proportions of high-sensitivity species, but intermediate levels of intact habitat protection (caramel in Fig. 2D-E, Fig. 3), also mainly concentrated in tropical ecoregions, whereas 72 have intermediate proportions of high-sensitivity species but low intact habitat protection, found not only in tropical eastern South America but also in North American temperate grasslands (salmon in Fig. 2D-E; Fig. 3).

Conversely, 29 ecoregions, mainly in the Amazonian basin (including the eastern slope of the Andes) have high proportion of high-sensitivity species while being relatively well covered by intact protected habitat (purple in Fig. 2D-E), mainly corresponding to tropical and subtropical moist broadleaf forests and mangroves (Fig. 3).

#### Across species

Among the 1,106 high-sensitivity species, 825 (75%) are inadequately covered by intact protected habitat, including 266 (24%) with no or only minor coverage. Among the later, 139 are threatened with extinction and thus of major conservation concern (Fig. 2C; Table S3). The latter are concentrated in the tropical Andes, Venezuela Coastal Range, and the Atlantic Forest of Brazil, but also in Central America, and the central plains of North America (Fig. 2C). When compared with all breeding species, they are disproportionately sedentary (98% *vs*. 83%), found in forest habitats (82% *vs*. 60%), specialised (average of 3.9 *vs*. 3.3), and have disproportionately small breeding ranges (average of 1.9×10^4^ *vs*. 1.9×10^6^). Nearly all (134 out of 139) of these species of major concern are actually in need of land protection because threatened by habitat loss (i.e., “Residential and commercial development” or “agriculture and aquaculture”; BirdLife International (2019)).

### Are protected areas retaining intact habitats over time?

Between 2000 and 2013, protected area intactness: decreased in 161 ecoregions, concentrated in South and Central America; was stable (i.e., |change| < 0.1) in 114 ecoregions, mostly in western North America and the Amazonian basin; and increased in 50 ecoregions, mainly in eastern North America (Fig. 2G). Trends in protected area intactness are negatively correlated with the proportion of high-sensitivity species per ecoregion (coef = – 0.22, P=6×10^−5^; Fig. 5B), with most ecoregions with >10% of local species being high-sensitivity (108 of 146; 74%) having experienced a decline in intactness (Fig. 2F).

## Discussion

We introduce here a new index of species’ sensitivity to anthropogenic land use changes (as measured through the human footprint), derived from field data on the variation of species’ abundance over their distributions. We focused on the breeding grounds, but the same methods can be applied to estimate sensitivity during other parts of the species’ annual cycle (which may also present important conservation challenges; e.g., Runge et al., 2015; Gaget et al., 2020) as species sensitivity probably varies with season. Even though adequate field data are not available for the vast majority of species in most regions, citizen science datasets (e.g. eBird, iNaturalist) are creating increasingly large datasets, brought together through data sharing platforms (e.g. GBIF), rendering it possible to estimate sensitivity for a growing number of species across the globe. The importance of measuring sensitivity directly from field records is stressed by the relatively poor link between sensitivity and traits (R^2^=0.18), suggesting that traits are a poor proxy for sensitivity, even though in our case results are robust as similar spatial distributions were obtained when using all *versus* only data-rich species (Fig. S6).

Breeding bird species in the Americas vary widely in their sensitivity to human pressure, as evidenced by the diversity of relationships between abundance and human footprint (Fig. 1). This supports previous results finding similarly wide variations (Rosenberg et al., 2019; Clavel et al., 2011; Guetté et al., 2017; Phalan et al., 2011; Williams et al., 2017). There is substantial overlap between threatened and high-sensitivity species (24.0% of high-sensitivity species are threatened, compared to 11.8% for all species), but the two measures are not the same. Indeed, whereas sensitivity to human footprint can be seen as an intrinsic ecological trait, threat levels result from the interaction between sensitivity and exposure to human pressure. We found that sensitivity is highly structured in space, being particularly dominant among species within tropical forest ecoregions, especially in the Andes, but also Central America and in the Amazonian basin (Fig. 2A), while being lower in temperate and boreal ecoregions. These results align with previous studies that reported a high sensitivity of tropical forest assemblages to even low levels of human pressure (Gibson et al., 2011; Barlow et al., 2016; Newbold et al., 2020).

These results confirm our prediction that the need for highly intact protected habitat is not the same everywhere, at least when it comes to the conservation of bird species. Placed in the context of the land sparing/land sharing debate (Green et al., 2005; Phalan et al., 2011; Williams et al., 2017), our results suggest that the best way for reconciling conservation and socio-economic targets varies across regions. A land sparing strategy is particularly crucial in the tropics, where bird assemblages include many species whose conservation depends on the maintenance of intact habitats, and that may thus require the establishment of strict protected areas. In contrast, land sharing may prove a more suitable strategy in temperate and boreal ecosystems, where fewer species depend on intact habitats, and can thus persist in areas with moderate levels of human activity, including multiple-use protected areas or even unprotected habitats. This said, we found high-sensitivity species in each one of the 325 ecoregions analysed (Table S2), which means that the protection of at least some intact habitat is crucial across all latitudes and biomes, in order to ensure the long-term persistence of all bird species.

Intact habitat protection – as quantified by the proportion of each ecoregion covered by intact protected habitat – is also widely variable (0% to 81%) and highly structured in space (Fig. 2B), being stronger in the Amazonian basin and some high latitude ecoregions (Boreal and Patagonian) that combine both low average human footprint and relatively high levels of protected area coverage. Worryingly, it does not correlate with the distribution of high-sensitivity species, the ones that need the protection of intact habitats the most. This is consistent with previous work that also found no correlation between the location of protected areas and species’ conservation needs, as measured by the presence of threatened species (instead, protected areas are mostly placed in areas of low economic interest; Venter et al., 2018). Of particular concern are the 58 ecoregions that have very low levels (<5%) of intact habitat protection despite hosting bird assemblages with a large proportion (>14%) of high-sensitivity species (Fig. 2D). These overlap extensively with Biodiversity Hotspots (i.e. biogeographic regions of exceptional plant endemism that have already lost >70% of their natural habitat, Mittermeier et al., 2004; Myers et al., 2000), particularly with the Tropical Andes, Tumbes-Chocó-Magdalena, Mesoamerica, and Cerrado hotspots (49 of the 58 ecoregions overlap one of these hotspots by >90%). Furthermore, many of these ecoregions (particularly in the Tropical Andes and Central America) cover areas identified as urgent priorities for the expansion of the global network of protected areas (Rodrigues et al., 2004; Butchart et al., 2015).

A complementary perspective is obtained by analysing mismatches between sensitivity and protected area coverage at the species level. This is particularly important as some narrow-range species, while living in ecoregions with low intact habitat protection, could be adequately covered if the distribution of intact protected habitats within an ecoregion matches species’ distribution. We identified 266 high-sensitivity species whose distributions have no or only minor coverage by intact protected habitat. The latter include species that are not protected at all (Maxwell et al., 2020) as well as others whose distributions are apparently well covered by protected areas but these are dominated by transformed habitats. For instance, the Critically Endangered Santa Marta wren, *Troglodytes monticola* (sensitivity = 41), is protected across 99.7% of its range, and would thus be considered as very adequately covered based on protected area coverage alone (Butchart et al., 2015), yet we found no intact protected habitat within its range. This corresponds to its Red List assessment, which highlights a long history of severe deforestation and degradation across the species range, which continues apace despite protection (IUCN, 2018). This example illustrates the importance of taking species’ sensitivity into account when evaluating the effectiveness of existing networks of protected areas as well as when identifying new priority areas for protection.

The persistence of many high-sensitivity species requires the establishment and effective management of strict protected areas, or other adequate mechanisms that guarantee the long-term maintenance of sufficient extents of intact habitats. This is all the most urgent for the 139 species we highlighted as being of major concern because they are simultaneously high-sensitivity, have no or only minor coverage by intact protected habitats, and are already at risk of extinction (Fig. 2C, Table S3). With nearly all (134; 96%) threatened by habitat loss, the mismatch between their high-sensitivity and the poor coverage of their range by intact protected habitats may prove dramatic in the near future in the absence of active measures to protect any remaining intact patches. The distributions of these species highlight ecoregions overlapping the same Biodiversity Hotspots as above, as well as the Atlantic Forest hotspot, all of which have already suffered major loss and transformation of their habitats (Williams et al., 2020). Restoration of currently degraded habitat is likely to play a key role in the long-term conservation of these species (Bull et al., 2020; Benayas et al., 2009), and indeed the regions where they occur are recognised global priorities for ecosystem restoration (Strassburg et al., 2020).

The importance of effective protected areas in these ecoregions will increase over time as pressures outside mount. Indeed, previous studies have shown that protected areas are becoming the last bastions for some species (Pacifici et al., 2020; Boakes et al., 2018). Unfortunately, though, we found that ecoregions with higher proportions of high-sensitivity species have experienced a faster degradation in the intactness of their protected areas, indicative of a growing mismatch between species needs and the availability of intact protected habitat (Fig. 2F-G). Previous studies had already raised stern warnings regarding the mounting human pressure within protected areas (Jones et al., 2018; Geldmann et al., 2014, 2019), through ongoing habitat loss and degradation (Spracklen et al., 2015; Cuenca et al., 2016; Bruner et al., 2001; Moore et al., 2017). Here we show that these trends are faster precisely where they are the most impactful: in regions where species need intact habitats the most.

Overall, our results show that the Americas’ protected area network is distributed such that it is not strategically located to conserve those bird species that need it the most, undermining its effectiveness in achieving the long-term conservation of nature (Rodrigues and Cazalis, 2020; SCBD, 2010), and highlight the importance of considering the habitat quality of protected areas (Barnes et al., 2018; Visconti et al., 2019). We highlight ecoregions and species where it is particularly urgent to ensure that remaining intact habitat is preserved, through protected areas and other relevant mechanisms, potentially complemented by the restoration of degraded habitat. With these ecoregions and species mostly concentrated in countries with limited economic resources, international cooperation is key to meeting this goal.

