## Supplementary materials for "Mismatch between bird species sensitivity and the protection of intact habitats across the Americas"

### Supplementary methods

#### 1) Alignment with BirdLife International taxonomy

The taxonomy followed by eBird (source of bird observation records) is different from the one followed by BirdLife International (source of species’ ranges, and Red List status). We aligned the former with the latter by: (1) replacing the species name used in eBird by the one in BirdLife International in simple cases of different names (but no taxonomic mismatch); (2) merging different eBird species (and summing their local abundance, if applicable) whenever they are lumped into a single species in the BirdLife International taxonomy; and (3) splitting single eBird species that correspond to multiple species in the BirdLife International taxonomy, using species distribution maps to separate the corresponding records (see Cazalis et al., (2020) for more details and Supplementary Spreadsheet 1 for list of species concerned by these changes).

#### 2) Calculating observer calibration index

We calculated an individual observer calibration index, closely related to the one calculated in Cazalis et al., (2020) (following Kelling et al., (2015) and Johnston et al., (2018)), for each observer included in the analyses (i.e., with more than a given threshold of experience: ≥ 50 checklists for a total of ≥ 100 species in the Americas during the study period). This index was calculated from a subset of the eBird records, selected through a somewhat different filtering process from the one used in the main analyses. As in the main analyses, we kept only checklists that reported all species detected, observations from 2010-2019, and excluding disapproved observations; in contrast, we did not restrict the dataset based on protocol type or sampling effort.

For any given observer, we calculated the observer calibration index as the log-scaled number of species an observer is expected to report on average during a standard sampling event. Usually this is done by running a mixed-effects model with the richness of each checklist as the response variable, with sampling effort and ecological drivers of species richness as explanatory variables, and with observer as a random effect (Johnston et al., 2018; Kelling et al., 2015; Cazalis et al., 2020). In this study, because of computing limitations, we split this calculation into two models: first we fitted a GAM assuming a negative binomial distribution (using function *bam* from the *'mgcv'* package (Wood, 2011)) with the following structure: species_richness ~ protocol_type + number_observers + s(duration) + s(starting_time) + te(longitude, latitude, Julian_day), with starting_time the time at which the sampling event started, s() a smoothed term enabling complex relation between variables, and te() an interacting smooth term enabling here richness to vary across space and time. We then extracted residuals of this model and used them in a linear mixed-model (using the function *lme* from the *'nlme'* package (Pinheiro et al., 2020)) with no explanatory variable and a random effect on observer. Finally, we simulated a sampling event under the “Stationary Points” protocol, fixing number of observers, duration, starting time, longitude, latitude and Julian day to their median values. We predicted the log-scaled species richness that should be detected in this hypothetical sampling event according to the GAM, and summed it with the random effect of each observer in order to get an observer calibration index per individual observer.

#### 3) Restricting bird records to breeding season and breeding grounds

We have restricted our dataset of bird records to observations made within each species’ broad breeding season and breeding grounds. Bird breeding season is known to vary with latitude (Baker, 1939). Although not all species within a given latitude necessarily breed at the same time (Baker, 1939), we did not have sufficiently detailed information to take these differences into account, and have instead assumed that within a given latitudinal band all species breed within the same broad season. To delimit the breeding season within each 10° latitudinal band, we studied the temporal distribution of the breeding codes that are sometimes associated with eBird observations (available for 1.2% of the observations in our dataset, N=1,532,862). First, we restricted the breeding codes to those that either correspond to probable breeding or confirmed breeding (i.e., by removing codes: “Flyover”, which does not correspond to a breeding behaviour; and “In appropriate habitat” and “Singing male”, which correspond to possible breeding only). We then plotted the temporal distribution of these breeding records (all species combined) per 10° latitudinal bands (Fig. S1). Based on these distributions, we considered that breeding occurs all year round between latitudes -10° and 10°. For other latitudinal bands, we defined the limits of the breeding season (Table S1) as the period containing 95% of the observations with breeding codes, bounded by circular quantiles at 2.5 and 97.5% of observations (black lines on Fig. S1) using the *circular* package (Agostinelli and Lund, 2017). For each species, in each latitudinal band, we included only records inside the boundaries of the breeding season.

In addition, we restricted the observations of each species to their respective breeding grounds, based on the BirdLife International’s distribution maps (BirdLife International and HBW, 2019), more specifically the polygons with Season codes 1 (resident) or 2 (breeding season only) and Presence codes 1 (extant), 2 (probably extant), 3 (possibly extant), and 4 (possibly extinct).

Within these temporal and spatial constraints, we considered each species as absent (abundance = 0) in all checklists where it was not detected and as present (with abundance as reported in the checklist) otherwise.

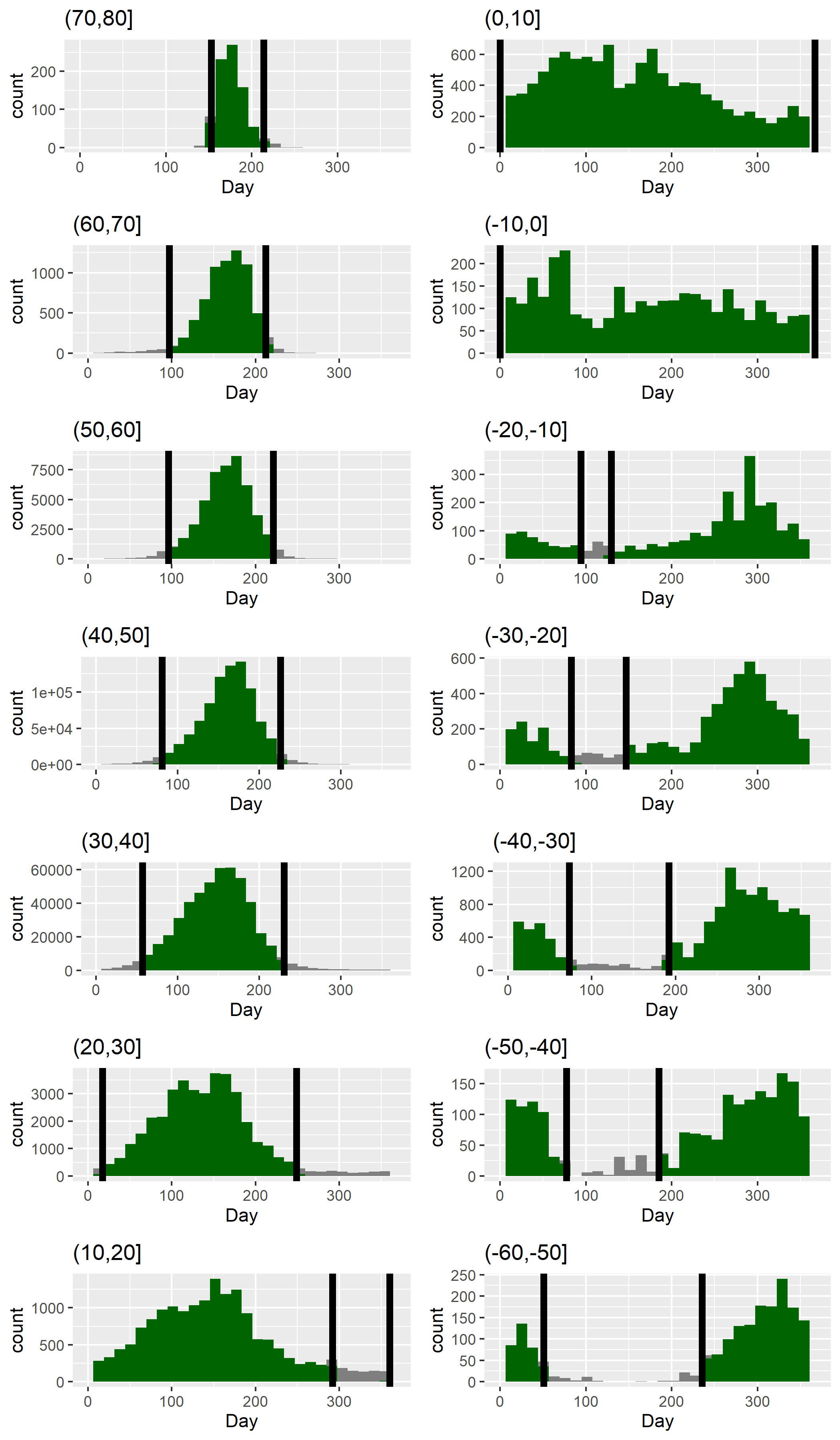

Fig. S1: Temporal distributions of eBird observations with associated breeding codes, per 10° latitudinal band. Vertical lines represent the limits of the breeding season as we defined it (i.e., the 2.5 and 97.5% circular quantiles; see Table S1). Bar colours show whether plotted data fall within (green) or outside (grey) the breeding season.

Table S1: Breeding season per latitudinal band. For two tropical bands (-10° to 0° and 0° to 10°), we considered that breeding occurs all year round. For all other bands, beginning and end dates were derived from the temporal distribution of breeding codes in eBird records (Fig. S1), as the 2.5 and 97.5% circular quantiles of Julian days.

|  | **Breeding season (Julian days)** | |
| --- | --- | --- |
| **Latitudinal band (degrees)** | **Beginning** | **End** |
| (70,80] | 153 | 214 |
| (60,70] | 97 | 212 |
| (50,60] | 96 | 221 |
| (40,50] | 81 | 226 |
| (30,40] | 57 | 230 |
| (20,30] | 17 | 249 |
| (10,20] | 360 | 292 |
| (0,10] | 0 | 367 |
| (-10,0] | 0 | 367 |
| (-20,-10] | 129 | 94 |
| (-30,-20] | 147 | 83 |
| (-40,-30] | 192 | 73 |
| (-50,-40] | 185 | 77 |
| (-60,-50] | 236 | 51 |

#### 4) Imputing species sensitivity

In order to estimate the sensitivity of the 1,874 data-poor species (i.e. those among the 4,424 that breed in the study area for which we were not able to measure sensitivity directly), we modelled the link between the sensitivity of the 2,550 data-rich species using the following linear model:

Sensitivity ~ Primary_Habitat + Primary_Diet + specialisation + log(Mass) + RedList + log(Range) + Migration + Order

Even though this model does not have a strong explaining power (R^2^=0.18), results indicate that major habitat significantly affects species sensitivity (P<10^-15^), with forest, grassland and riparian species being particularly sensitive, while species favouring artificial habitats, deserts, and savannahs being on average less sensitive to human footprint (Fig. S2A). Taxonomic Order also greatly influences species sensitivity (P=3.10^-8^), with Rheiformes, Phoenicopteriformes, and Cariamiformes at highest sensitivity, while Psittaciformes, Podicipediformes, Falconiformes, Columbiformes, Apodiformes, and Accipitriformes showed lower sensitivity (Fig. S2B). Species diet had a slightly significant effect on sensitivity (P=0.027) with high sensitivity for nectarivorous and low sensitivity for granivorous (Fig. S2C). Migration status greatly influenced sensitivity, with sedentary species showing higher sensitivity than strict migrating species than partial migrating species (P=9.10^-6^; Fig. S2D). Species sensitivity increased with species specialisation (P<10^-15^, Fig. S2F), decreased with species range size (log-scaled) in the Americas (P=0.049, Fig. S2G), increased with species body mass (P=4.10^-4^, Fig. S2H), and increased with species quantitative Red List status (P=0.049, Fig. S2E).

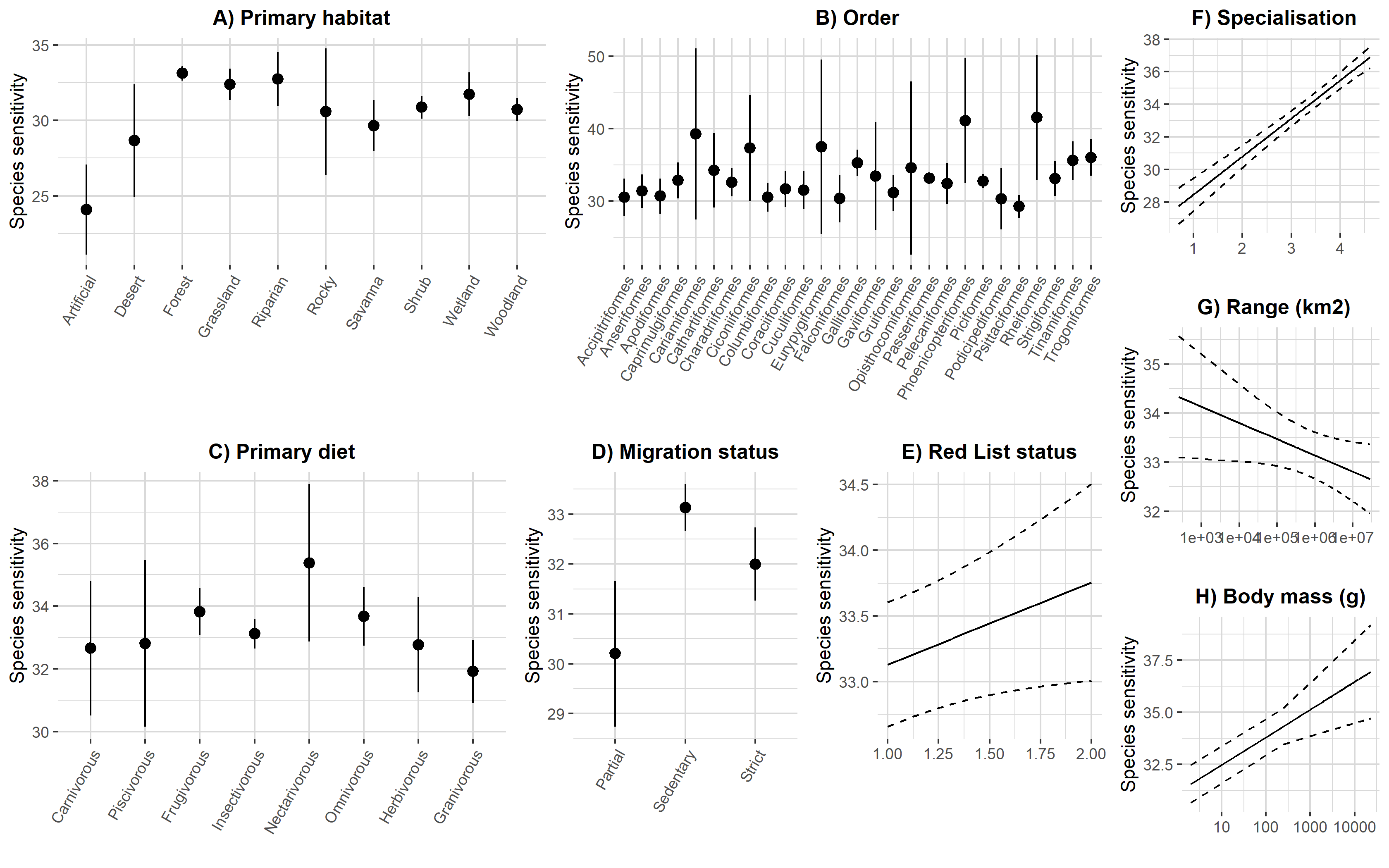

Fig. S2: Effects of species traits on sensitivity to human footprint, for 8 traits: (A) primary habitat, (B) taxonomic Order, (C) primary diet, (D) migration status, (E) Red List status, (F) habitat specialisation, (G) range size, and (H) body mass.

Using the estimates of this model, we then imputed sensitivity for the 1,874 data-poor bird species based on their traits and using the R function *predict*. Two species belong to orders that were not represented within the 2,550 data-rich species for which we measured sensitivity directly (respectively Bucerotiformes and Pterocliformes), we thus assigned to these species the sensitivity value of the Order for which sensitivity was median (Caprimulgiformes).

This first estimate of imputed sensitivity was biased (as we can see when comparing the measured sensitivity for the 2,550 data-rich species with the estimate that would arise from model predictions; Fig. S3A). We thus corrected it, by scaling the imputed sensitivity and then reversing the scaling using the measured sensitivity parameters (i.e., multiplying by the standard deviation of measured sensitivity and adding its mean value). We then replaced the few imputed values below 0 by 0 and the few values above 50 by 50. This corrected the bias found in the first estimate (Fig. S3B).

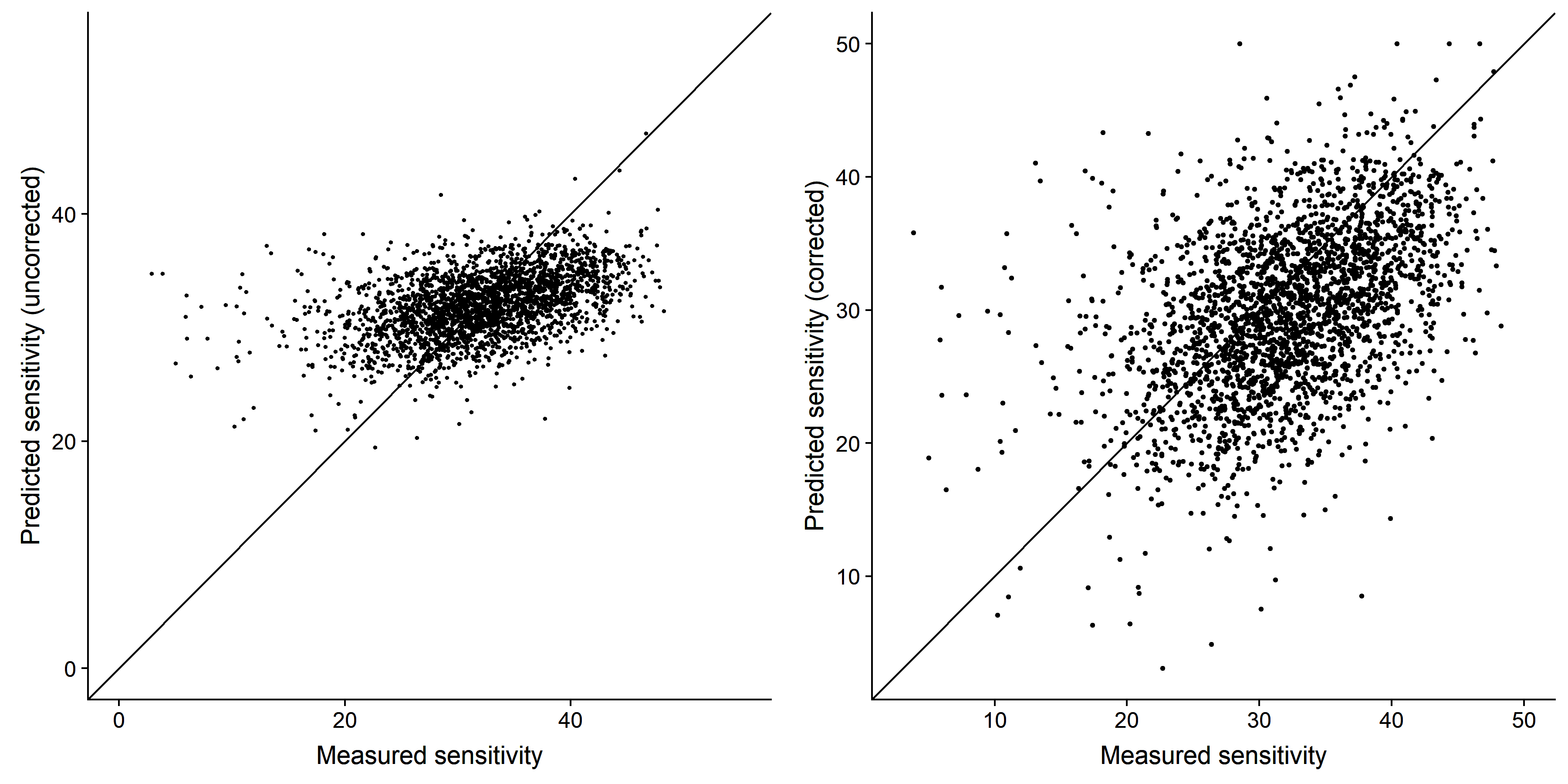

Fig. S3: Imputed sensitivity (left) and corrected imputed sensitivity (right) of the 2,550 species for which sensitivity was directly measured. Correction consisted in scaling the imputed sensitivity, multiplying by the standard deviation of measured sensitivity, adding the mean value of measured sensitivity and finally replacing values below 0 by 0 and values above 50 by 50.

We then used measured sensitivity for the 2,550 data-rich species and corrected imputed sensitivity for the 1,874 data-poor species (see both distributions in Fig. 1).

#### 5) Sampling bias towards sites with high human footprint

There was across the entire study area a strong bias in sampling towards sites with high human footprint and a near-complete absence of sampling in sites with very low human footprint (Fig. S4). For this reason, values of species’ sensitivity cannot be interpreted in absolute terms: a sensitivity of 30 does not (necessarily) mean that 30% of the population is found in areas of human footprint lower than 20 (= 50-30). Instead, we use values of species’ sensitivity in relative terms, to compare sensitivity of species and of regions.

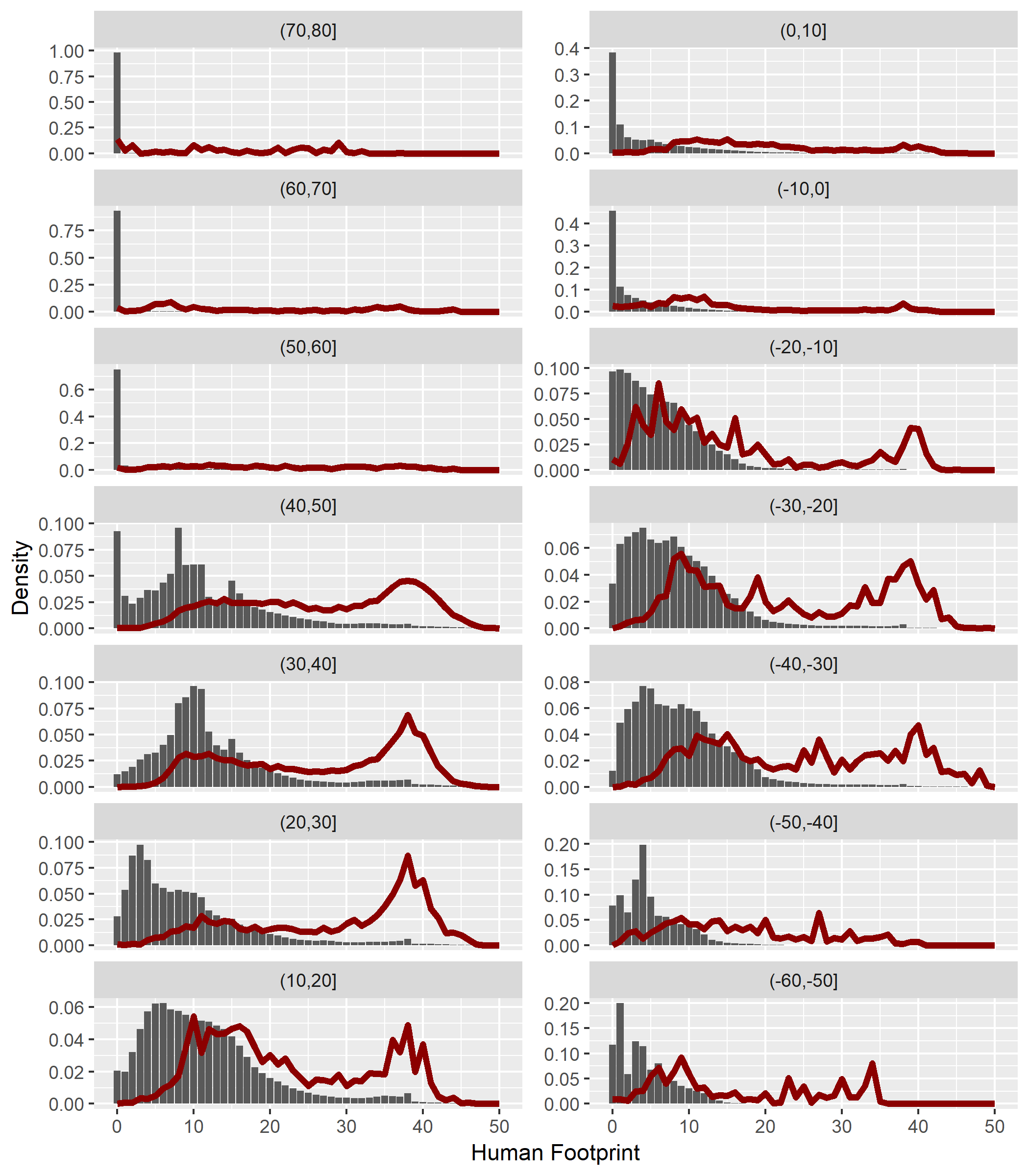
 Fig. S4: Distribution of human footprint raster cells (grey bars) compared with distribution of human footprint of checklists (red lines) per 10° latitudinal bands in the study area.

### Supplementary results

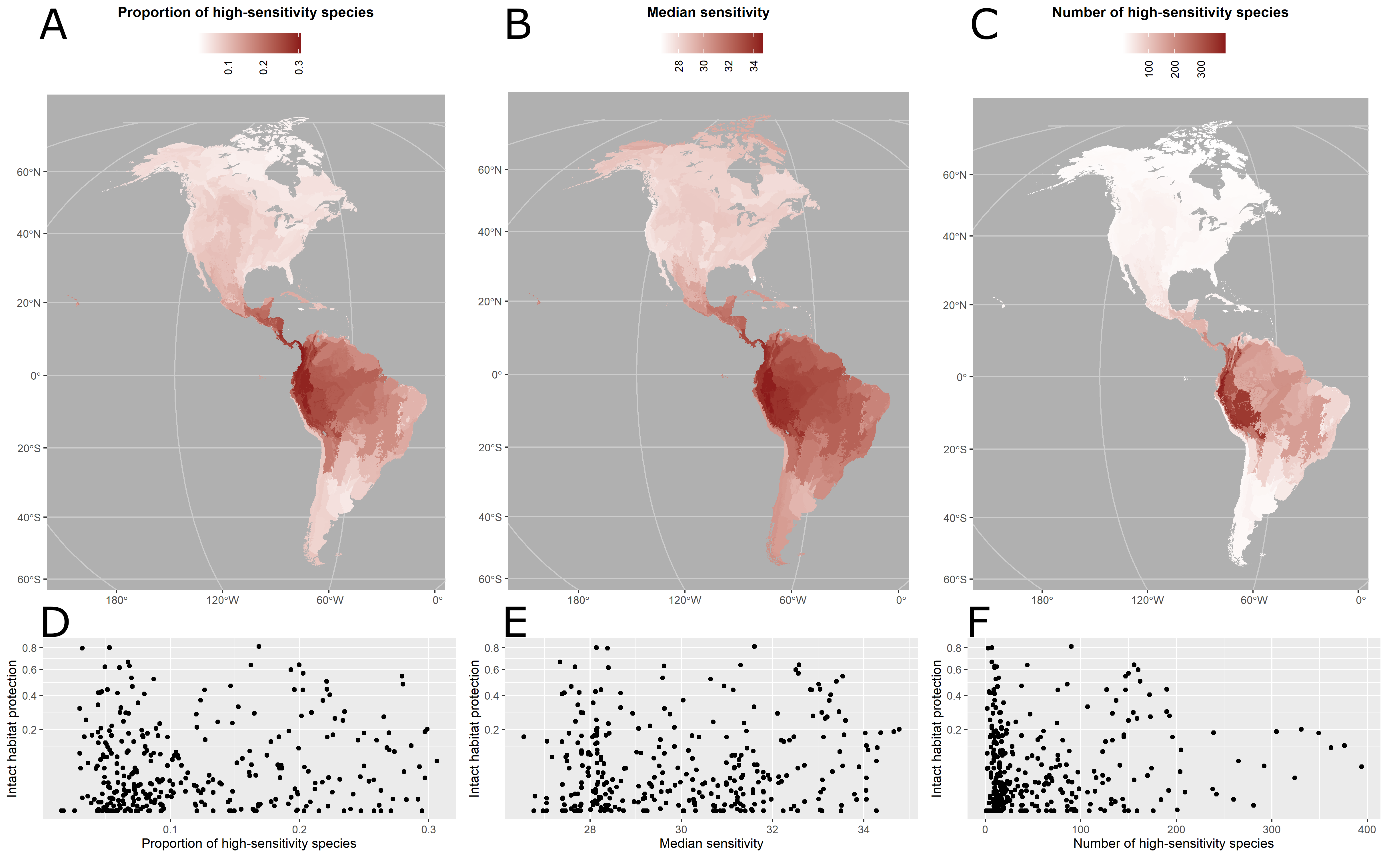

Fig. S5: Spatial patterns in three ecoregion-level metrics of the sensitivity of bird communities to human footprint: (A) median sensitivity across all species that breed in the ecoregion; (B) number of high-sensitivity species that breed in the ecoregion; and (C) proportion of all species breeding in the ecoregion that are high-sensitivity species and correlation with intact habitat protection (D-F). Intact habitat protection does not correlate with proportion of high-sensitivity species (coef = 0.044, P=0.424; D), with median sensitivity (coef = 0.071, P=0.199; E), or with number of high-sensitivity species (coef = 0.058, P=0.297; F).

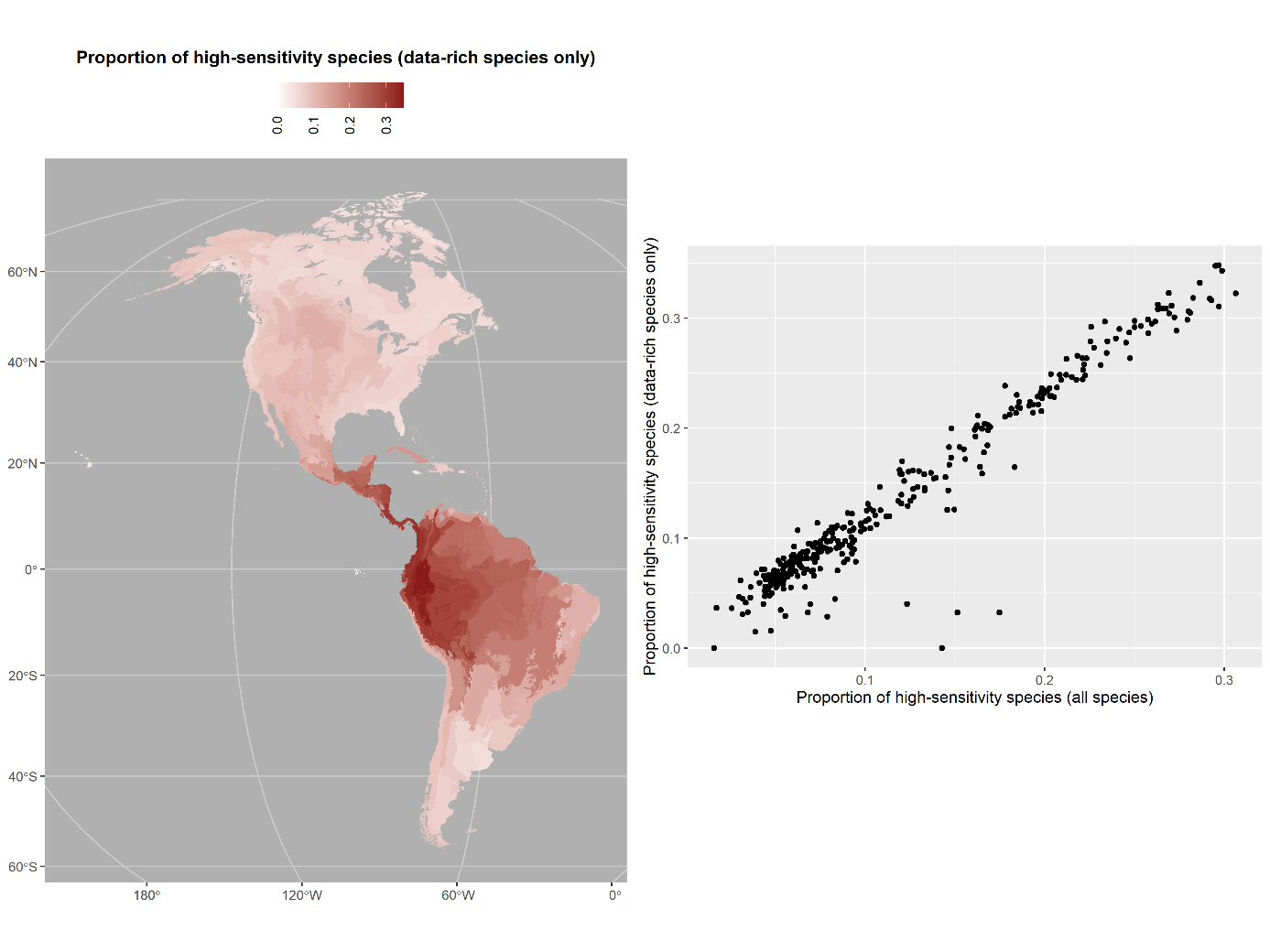

Fig. S6: Proportion of high-sensitivity species using only data-rich species (for which we were able to measure sensitivity directly; N=2,550, of which 525 are high-sensitivity species): (A) distribution across ecoregions; (B) correlation with the proportion of high-sensitivity species based on all species (N=4,424, of which 581 are high sensitivity species; coef = 0.969, P<2.10^-16^).

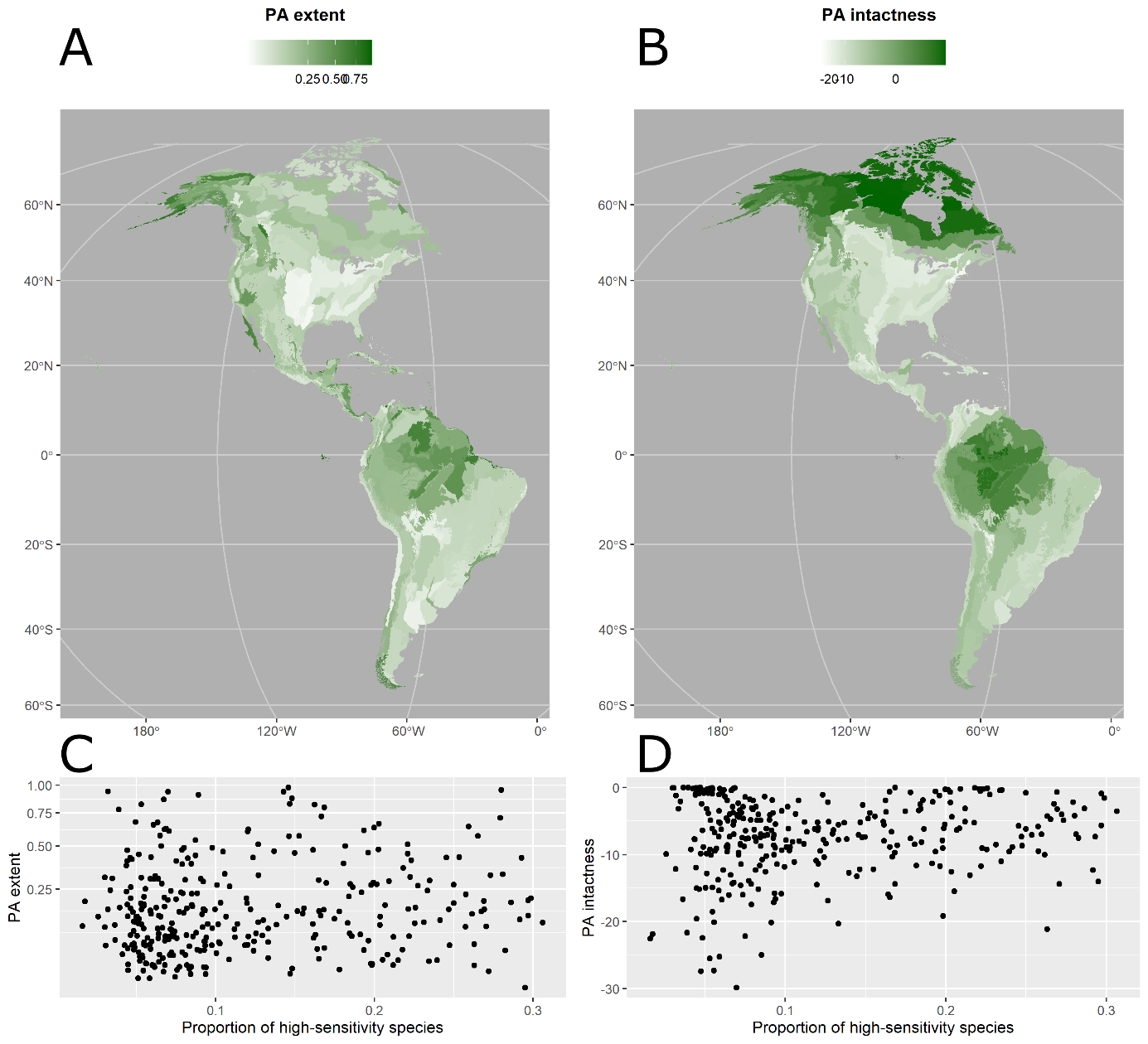

Fig. S7: Spatial patterns of the investment in habitat protection across ecoregions, as measured by: (A) protected area extent (fraction of the ecoregion covered by protected areas), and (B) protected area intactness (mean intactness of pixels within the ecoregion that intersect by >50% with protected areas), and the respective (C, D) correlation with the proportion of high-sensitivity species.

Table S2: Results at the ecoregion level for the 325 ecoregions used in the study.

| **Ecoregion**  **name** | **Eco.**  **code** | **Extent** | **PA**  **extent** | **Intact habitat protection** | **PA intactness** | **Trend in PA intactness** | **Proportion of high-sensitivy**  **species** | **Number of high-sensitivity species** | **Median sensitivity of species** |
| --- | --- | --- | --- | --- | --- | --- | --- | --- | --- |
| Alaska Peninsula montane taiga | 7 | 4.79E+10 | 0.83 | 0.8 | -0.3 | 0.04 | 0.05 | 7 | 28.14 |
| Alaska/St. Elias Range tundra | 8 | 1.52E+11 | 0.45 | 0.44 | -0.3 | 0 | 0.05 | 9 | 28.24 |
| Alberta Mountain forests | 10 | 3.99E+10 | 0.68 | 0.62 | -0.8 | 0.02 | 0.05 | 12 | 27.66 |
| Alberta/British Columbia foothills forests | 11 | 1.21E+11 | 0.02 | 0.01 | -4.9 | 0.48 | 0.06 | 17 | 28.12 |
| Aleutian Islands tundra | 14 | 5.48E+09 | 0.94 | 0.79 | -1.2 | 0.17 | 0.03 | 3 | 28.38 |
| Allegheny Highlands forests | 15 | 8.41E+10 | 0.04 | 0 | -9.2 | 0.93 | 0.04 | 8 | 28.19 |
| Alta Parano Atlantic forests | 17 | 4.85E+11 | 0.06 | 0.01 | -9.4 | -0.31 | 0.1 | 81 | 31.2 |
| Alvarado mangroves | 21 | 4.56E+09 | 0.28 | 0.03 | -8.2 | -0.1 | 0.18 | 64 | 31.32 |
| Amapa mangroves | 22 | 1.57E+09 | 0.8 | 0.81 | 0 | 0 | 0.17 | 90 | 31.6 |
| Appalachian mixed mesophytic forests | 32 | 1.93E+11 | 0.03 | 0 | -10.1 | 0.92 | 0.05 | 11 | 28.15 |
| Appalachian/Blue Ridge forests | 33 | 1.60E+11 | 0.06 | 0.01 | -7.3 | 0.54 | 0.06 | 12 | 28.12 |
| Apure-Villavicencio dry forests | 35 | 6.87E+10 | 0.11 | 0.01 | -9.3 | -0.57 | 0.23 | 209 | 33.06 |
| Araucaria moist forests | 38 | 2.17E+11 | 0.04 | 0.01 | -7.8 | -0.37 | 0.1 | 61 | 31.09 |
| Araya and Paria xeric scrub | 39 | 5.29E+09 | 0.05 | 0 | -15.8 | -1.1 | 0.09 | 39 | 30.06 |
| Arctic coastal tundra | 40 | 9.84E+10 | 0.13 | 0.12 | -0.2 | -0.01 | 0.05 | 6 | 29.51 |
| Arctic foothills tundra | 42 | 1.29E+11 | 0.21 | 0.22 | -0.2 | -0.02 | 0.05 | 7 | 29.32 |
| Argentine Espinal | 43 | 1.09E+11 | 0.02 | 0 | -6.7 | -0.27 | 0.06 | 17 | 29.16 |
| Argentine Monte | 44 | 4.10E+11 | 0.08 | 0.06 | -3.7 | -0.15 | 0.07 | 24 | 29.92 |
| Arid Chaco | 45 | 9.91E+10 | 0.03 | 0.02 | -3.9 | -0.07 | 0.05 | 16 | 29.35 |
| Arizona Mountains forests | 46 | 1.09E+11 | 0.1 | 0.06 | -3.5 | 0.05 | 0.1 | 26 | 27.83 |
| Aruba-Curacao-Bonaire cactus scrub | 48 | 4.60E+08 | 0.1 | 0 | -22.5 | -0.92 | 0.02 | 1 | 27.05 |
| Atacama desert | 50 | 1.05E+11 | 0.02 | 0.01 | -4.8 | -0.55 | 0.08 | 12 | 29.19 |
| Atlantic Coast restingas | 51 | 7.89E+09 | 0.16 | 0.03 | -11.8 | -0.74 | 0.1 | 70 | 31.16 |
| Atlantic coastal pine barrens | 54 | 8.98E+09 | 0.26 | 0 | -15.1 | 3.03 | 0.05 | 8 | 28.06 |
| Atlantic dry forests | 55 | 1.15E+11 | 0.07 | 0.05 | -2.8 | -0.06 | 0.07 | 35 | 30.33 |
| Baffin coastal tundra | 61 | 9.12E+09 | 0.05 | 0.05 | -0.2 | -0.16 | 0.04 | 2 | 28.75 |
| Bahamian dry forests | 62 | 4.81E+09 | 0.01 | 0 | -11.6 | -0.36 | 0.08 | 9 | 27.73 |
| Bahamian mangroves | 63 | 6.63E+09 | 0.02 | 0 | -7.5 | -0.01 | 0.09 | 10 | 27.8 |
| Bahia coastal forests | 65 | 1.10E+11 | 0.08 | 0.01 | -8.3 | -0.37 | 0.13 | 79 | 31.57 |
| Bahia interior forests | 66 | 2.30E+11 | 0.04 | 0.01 | -7.8 | -0.35 | 0.12 | 91 | 31.57 |
| Bahia mangroves | 67 | 2.12E+09 | 0.19 | 0.09 | -8.8 | -0.68 | 0.1 | 49 | 31.11 |
| Baja California desert | 68 | 7.79E+10 | 0.61 | 0.46 | -2.3 | -0.11 | 0.07 | 13 | 27.58 |
| Bajio dry forests | 69 | 3.75E+10 | 0.07 | 0.02 | -10.7 | -0.1 | 0.09 | 25 | 29.33 |
| Balsas dry forests | 71 | 6.26E+10 | 0.11 | 0.02 | -7.7 | 0.03 | 0.14 | 56 | 30.52 |
| Belizean Coast mangroves | 75 | 2.80E+09 | 0.11 | 0 | -12.6 | -1 | 0.2 | 69 | 31.81 |
| Belizian pine forests | 77 | 2.84E+09 | 0.03 | 0 | -6.1 | -0.77 | 0.21 | 69 | 31.8 |
| Beni savanna | 78 | 1.26E+11 | 0.01 | 0 | -7.6 | -0.07 | 0.18 | 136 | 32.7 |
| Beringia lowland tundra | 80 | 1.51E+11 | 0.66 | 0.62 | -0.5 | -0.02 | 0.06 | 10 | 28.4 |
| Beringia upland tundra | 81 | 9.76E+10 | 0.43 | 0.42 | -0.1 | 0 | 0.06 | 9 | 28.67 |
| Blue Mountains forests | 84 | 6.48E+10 | 0.09 | 0.04 | -4.4 | 0.01 | 0.07 | 16 | 28.17 |
| Bocas del Toro-San Bastimentos Island-San Blas mangroves | 85 | 5.42E+08 | 0.23 | 0 | -9.5 | -0.49 | 0.26 | 109 | 32.56 |
| Bolivian montane dry forests | 88 | 8.05E+10 | 0.02 | 0 | -19.2 | -1.2 | 0.2 | 201 | 33.04 |
| Bolivian Yungas | 87 | 9.08E+10 | 0.02 | 0.01 | -3.1 | -0.13 | 0.21 | 238 | 33.12 |
| British Columbia mainland coastal forests | 94 | 1.38E+11 | 0.3 | 0.29 | -0.7 | -0.04 | 0.06 | 16 | 27.8 |
| Brooks/British Range tundra | 95 | 1.60E+11 | 0.63 | 0.63 | 0 | 0 | 0.07 | 10 | 29.62 |
| Caatinga | 97 | 7.36E+11 | 0.07 | 0.03 | -5.1 | -0.22 | 0.11 | 69 | 31.07 |
| Caatinga Enclaves moist forests | 98 | 4.81E+09 | 0.28 | 0.01 | -13 | -1.76 | 0.07 | 21 | 29.56 |
| California Central Valley grasslands | 100 | 5.52E+10 | 0.04 | 0 | -12.8 | 0.23 | 0.05 | 12 | 27.43 |
| California coastal sage and chaparral | 101 | 3.63E+10 | 0.09 | 0.02 | -8.5 | 0.24 | 0.07 | 17 | 27.39 |
| California interior chaparral and woodlands | 102 | 6.47E+10 | 0.07 | 0.01 | -9.6 | 0.53 | 0.06 | 16 | 27.59 |
| California montane chaparral and woodlands | 103 | 2.05E+10 | 0.24 | 0.14 | -3.4 | 0.21 | 0.07 | 17 | 27.38 |
| Campos Rupestres montane savanna | 105 | 2.65E+10 | 0.26 | 0.06 | -8 | -0.26 | 0.11 | 78 | 31.26 |
| Canadian Aspen forests and parklands | 106 | 3.98E+11 | 0.08 | 0.01 | -11.4 | 0.38 | 0.07 | 22 | 28.43 |
| Caqueta moist forests | 112 | 1.85E+11 | 0.27 | 0.26 | -1 | -0.24 | 0.23 | 159 | 33.16 |
| Cascade Mountains leeward forests | 118 | 4.64E+10 | 0.41 | 0.35 | -1.4 | 0.02 | 0.08 | 19 | 28.07 |
| Catatumbo moist forests | 121 | 2.29E+10 | 0.28 | 0 | -8.6 | -0.27 | 0.17 | 110 | 31.61 |
| Cauca Valley dry forests | 122 | 7.36E+09 | 0.02 | 0 | -21.1 | 0.37 | 0.26 | 176 | 33.5 |
| Cauca Valley montane forests | 123 | 3.21E+10 | 0.13 | 0 | -12.3 | -0.3 | 0.29 | 260 | 34.06 |
| Cayman Islands dry forests | 125 | 1.34E+08 | 0.08 | 0 | -16 | 0 | 0.07 | 5 | 28.23 |
| Cayos Miskitos-San Andr<U+FFFD>s & Providencia moist forests | 126 | 9.54E+07 | 0.38 | 0 | -7.7 | -1 | 0.04 | 2 | 27.66 |
| Central American Atlantic moist forests | 130 | 8.97E+10 | 0.42 | 0.15 | -5.7 | -0.57 | 0.25 | 133 | 32.19 |
| Central American dry forests | 131 | 6.82E+10 | 0.08 | 0 | -11.3 | -0.42 | 0.19 | 132 | 31.3 |
| Central American montane forests | 132 | 1.33E+10 | 0.31 | 0.03 | -9 | -0.64 | 0.22 | 116 | 31.81 |
| Central American pine-oak forests | 133 | 1.12E+11 | 0.12 | 0.03 | -8.5 | -0.52 | 0.23 | 129 | 31.94 |
| Central and Southern Cascades forests | 161 | 4.50E+10 | 0.21 | 0.11 | -3.8 | 0.06 | 0.07 | 16 | 27.81 |
| Central and Southern mixed grasslands | 162 | 2.83E+11 | 0.01 | 0 | -8.7 | 0.17 | 0.07 | 18 | 27.79 |
| Central Andean dry puna | 136 | 3.08E+11 | 0.11 | 0.06 | -3.4 | -0.06 | 0.18 | 59 | 31.66 |
| Central Andean puna | 137 | 1.62E+11 | 0.14 | 0.04 | -6.6 | -0.07 | 0.17 | 123 | 31.9 |
| Central Andean wet puna | 138 | 1.18E+11 | 0.12 | 0.01 | -8.9 | -0.15 | 0.24 | 242 | 33.8 |
| Central British Columbia Mountain forests | 142 | 7.19E+10 | 0.09 | 0.09 | -0.4 | 0 | 0.05 | 10 | 28 |
| Central Canadian Shield forests | 143 | 4.63E+11 | 0.13 | 0.14 | -0.5 | 0 | 0.05 | 11 | 27.73 |
| Central forest/grasslands transition zone | 163 | 4.08E+11 | 0.02 | 0 | -13.9 | 1.41 | 0.08 | 19 | 28.35 |
| Central Mexican matorral | 150 | 5.95E+10 | 0.05 | 0.01 | -15.8 | -0.2 | 0.1 | 29 | 29.36 |
| Central Mexican wetlands | 151 | 2.80E+08 | 0.17 | 0 | -25 | 0.36 | 0.09 | 22 | 28.96 |
| Central Pacific coastal forests | 152 | 7.38E+10 | 0.12 | 0.09 | -2.4 | 0.04 | 0.05 | 10 | 27.16 |
| Central tall grasslands | 164 | 2.49E+11 | 0.02 | 0 | -14.2 | 1.36 | 0.07 | 14 | 28.64 |
| Central U.S. hardwood forests | 159 | 2.97E+11 | 0.03 | 0 | -10 | 0.89 | 0.07 | 13 | 28.06 |
| Cerrado | 165 | 1.92E+12 | 0.07 | 0.04 | -5.6 | -0.26 | 0.16 | 170 | 32.31 |
| Chaco | 166 | 6.11E+11 | 0.11 | 0.08 | -2.8 | -0.17 | 0.1 | 76 | 31.26 |
| Chiapas Depression dry forests | 174 | 1.41E+10 | 0.03 | 0 | -15.5 | 0.46 | 0.21 | 86 | 31.1 |
| Chiapas montane forests | 175 | 5.79E+09 | 0.04 | 0.02 | -5.2 | -0.47 | 0.22 | 91 | 31.69 |
| Chihuahuan desert | 176 | 5.11E+11 | 0.07 | 0.03 | -5.4 | -0.06 | 0.09 | 33 | 28.49 |
| Chilean matorral | 177 | 1.49E+11 | 0.01 | 0 | -7.6 | -0.28 | 0.08 | 13 | 30.27 |
| Chimalapas montane forests | 178 | 2.09E+09 | 0.14 | 0.06 | -4.4 | -0.09 | 0.2 | 74 | 31.16 |
| Chiquitano dry forests | 180 | 2.31E+11 | 0.01 | 0.01 | -2.6 | -0.1 | 0.15 | 115 | 31.79 |
| Choco-Darien moist forests | 181 | 7.38E+10 | 0.12 | 0.07 | -3.5 | -0.29 | 0.31 | 265 | 34.28 |
| Coastal Venezuelan mangroves | 184 | 5.86E+09 | 0.27 | 0.01 | -8.9 | -0.38 | 0.1 | 54 | 30.46 |
| Colorado Plateau shrublands | 186 | 3.27E+11 | 0.1 | 0.02 | -6.4 | 0.01 | 0.09 | 26 | 28.18 |
| Colorado Rockies forests | 187 | 1.33E+11 | 0.15 | 0.1 | -3.2 | 0.03 | 0.08 | 21 | 28.15 |
| Cook Inlet taiga | 189 | 2.79E+10 | 0.31 | 0.25 | -2 | -0.13 | 0.03 | 5 | 28.14 |
| Copper Plateau taiga | 192 | 1.72E+10 | 0.21 | 0.21 | -0.9 | 0.03 | 0.05 | 7 | 28.15 |
| Cordillera Central paramo | 193 | 1.22E+10 | 0.16 | 0.11 | -3.2 | -0.07 | 0.27 | 205 | 34.1 |
| Cordillera de Merida paramo | 196 | 2.82E+09 | 0.72 | 0 | -10.3 | -0.27 | 0.17 | 95 | 31.8 |
| Cordillera La Costa montane forests | 194 | 1.44E+10 | 0.57 | 0 | -12.2 | -0.84 | 0.15 | 85 | 31.23 |
| Cordillera Oriental montane forests | 195 | 6.80E+10 | 0.27 | 0.06 | -6.2 | -0.31 | 0.25 | 292 | 33.54 |
| Cordoba montane savanna | 209 | 5.83E+10 | 0.06 | 0.02 | -6.4 | -0.25 | 0.05 | 17 | 29.58 |
| Costa Rican seasonal moist forests | 198 | 1.07E+10 | 0.1 | 0 | -14.4 | -0.55 | 0.27 | 150 | 33.09 |
| Cuban cactus scrub | 203 | 3.27E+09 | 0.19 | 0 | -14.5 | -0.48 | 0.12 | 17 | 29.94 |
| Cuban dry forests | 204 | 6.59E+10 | 0.06 | 0 | -10.4 | -0.04 | 0.12 | 19 | 30.28 |
| Cuban moist forests | 205 | 2.14E+10 | 0.21 | 0 | -10.9 | -0.09 | 0.12 | 18 | 30.28 |
| Cuban pine forests | 206 | 6.44E+09 | 0.07 | 0 | -11.4 | 0.01 | 0.12 | 17 | 30.03 |
| Cuban wetlands | 207 | 5.68E+09 | 0.62 | 0.21 | -6 | -0.04 | 0.12 | 17 | 29.85 |
| Davis Highlands tundra | 213 | 8.81E+10 | 0.31 | 0.31 | -0.1 | -0.01 | 0.03 | 2 | 29.63 |
| East Central Texas forests | 223 | 5.28E+10 | 0.01 | 0 | -9.8 | 0.09 | 0.05 | 10 | 28.01 |
| Eastern Canadian forests | 235 | 4.88E+11 | 0.17 | 0.16 | -0.6 | 0.03 | 0.05 | 10 | 28.05 |
| Eastern Canadian Shield taiga | 234 | 7.55E+11 | 0.1 | 0.1 | 0 | 0 | 0.05 | 8 | 27.81 |
| Eastern Cascades forests | 236 | 5.53E+10 | 0.06 | 0.01 | -7 | 0.09 | 0.07 | 17 | 28.06 |
| Eastern Cordillera real montane forests | 238 | 1.03E+11 | 0.19 | 0.13 | -3.1 | -0.39 | 0.29 | 376 | 34.06 |
| Eastern forest/boreal transition | 252 | 3.48E+11 | 0.11 | 0.09 | -3.1 | 0.12 | 0.05 | 12 | 28.23 |
| Eastern Great Lakes lowland forests | 240 | 1.17E+11 | 0.02 | 0 | -16.7 | 1.74 | 0.06 | 14 | 28.4 |
| Eastern Panamanian montane forests | 250 | 3.05E+09 | 0.71 | 0.54 | -2.2 | -0.25 | 0.28 | 147 | 33.54 |
| Ecuadorian dry forests | 254 | 2.13E+10 | 0.03 | 0 | -9.5 | -0.17 | 0.24 | 121 | 32.4 |
| Edwards Plateau savanna | 255 | 6.19E+10 | 0.01 | 0 | -14.3 | -0.65 | 0.07 | 12 | 27.59 |
| Enriquillo wetlands | 260 | 6.32E+08 | 0.62 | 0.29 | -5.6 | -0.1 | 0.07 | 8 | 28.93 |
| Esmeraldes/Choco mangroves | 262 | 6.53E+09 | 0.32 | 0.11 | -4.9 | -0.46 | 0.27 | 123 | 33.47 |
| Everglades | 270 | 2.01E+10 | 0.25 | 0.16 | -3.2 | 0.1 | 0.03 | 5 | 26.55 |
| Flint Hills tall grasslands | 277 | 2.97E+10 | 0.02 | 0 | -9.4 | 0.26 | 0.06 | 9 | 27.67 |
| Florida sand pine scrub | 278 | 3.89E+09 | 0.16 | 0 | -12.1 | 0.28 | 0.03 | 5 | 27.4 |
| Fraser Plateau and Basin complex | 279 | 1.37E+11 | 0.1 | 0.1 | -0.4 | 0 | 0.07 | 16 | 27.97 |
| Great Basin montane forests | 288 | 5.80E+09 | 0.5 | 0.29 | -3.9 | 0.03 | 0.07 | 15 | 27.84 |
| Great Basin shrub steppe | 289 | 3.37E+11 | 0.09 | 0.04 | -4.6 | 0 | 0.07 | 20 | 27.95 |
| Greater Antilles mangroves | 293 | 1.07E+10 | 0.46 | 0.1 | -8.1 | 0.01 | 0.1 | 28 | 29.02 |
| Guajira-Barranquilla xeric scrub | 295 | 3.17E+10 | 0.04 | 0 | -15.9 | -0.78 | 0.16 | 93 | 31.23 |
| Guayanan Highlands moist forests | 296 | 3.38E+11 | 0.61 | 0.6 | -0.5 | -0.14 | 0.19 | 160 | 32.51 |
| Guayaquil flooded grasslands | 297 | 2.94E+09 | 0.02 | 0 | -7.9 | 0.14 | 0.2 | 72 | 31.4 |
| Guianan Freshwater swamp forests | 298 | 7.74E+09 | 0.08 | 0.06 | -3.6 | -0.27 | 0.16 | 85 | 31.6 |
| Guianan mangroves | 299 | 1.46E+10 | 0.57 | 0.32 | -3.8 | -0.69 | 0.15 | 107 | 31.6 |
| Guianan moist forests | 300 | 5.14E+11 | 0.29 | 0.29 | -0.6 | -0.16 | 0.17 | 133 | 32.11 |
| Gulf of California xeric scrub | 304 | 2.36E+10 | 0.49 | 0.41 | -1.9 | -0.12 | 0.08 | 10 | 27.38 |
| Gulf of Fonseca mangroves | 305 | 1.63E+09 | 0.67 | 0 | -8.8 | -0.36 | 0.06 | 13 | 28.71 |
| Gulf of Guayaquil-Tumbes mangroves | 306 | 3.32E+09 | 0.16 | 0 | -9.7 | -0.74 | 0.17 | 54 | 30.74 |
| Gulf of Panama mangroves | 308 | 2.43E+09 | 0.05 | 0 | -8.7 | -0.39 | 0.25 | 125 | 32.55 |
| Gulf of St. Lawrence lowland forests | 309 | 3.95E+10 | 0.03 | 0.02 | -4.9 | 0.45 | 0.05 | 10 | 27.81 |
| Gurupa varzea | 310 | 9.95E+09 | 0.46 | 0.44 | -1.2 | -0.38 | 0.2 | 127 | 32.83 |
| Guyanan savanna | 311 | 1.05E+11 | 0.22 | 0.16 | -2.5 | -0.49 | 0.18 | 145 | 32.46 |
| Hawaii tropical dry forests | 314 | 6.65E+09 | 0.16 | 0.07 | -4.6 | -0.38 | 0.15 | 15 | 30.98 |
| Hawaii tropical high shrublands | 315 | 1.86E+09 | 0.43 | 0.37 | -1.1 | 0 | 0.12 | 10 | 30.04 |
| Hawaii tropical low shrublands | 316 | 1.53E+09 | 0.08 | 0.03 | -5 | 0 | 0.07 | 6 | 29.41 |
| Hawaii tropical moist forests | 317 | 6.75E+09 | 0.13 | 0.09 | -3.5 | -0.19 | 0.17 | 18 | 31.17 |
| High Arctic tundra | 320 | 4.65E+11 | 0.1 | 0.1 | 0 | -0.02 | 0.03 | 2 | 29.63 |
| Hispaniolan dry forests | 325 | 1.55E+10 | 0.23 | 0 | -10.2 | -0.71 | 0.09 | 12 | 29.41 |
| Hispaniolan moist forests | 326 | 4.61E+10 | 0.11 | 0 | -11.4 | -0.17 | 0.09 | 13 | 29.54 |
| Hispaniolan pine forests | 327 | 1.16E+10 | 0.39 | 0.02 | -7.4 | -0.16 | 0.08 | 11 | 29.3 |
| Humid Chaco | 333 | 3.36E+11 | 0.07 | 0.02 | -6.4 | -0.18 | 0.08 | 43 | 30.8 |
| Humid Pampas | 334 | 2.41E+11 | 0.05 | 0 | -11.9 | -0.26 | 0.04 | 13 | 29.06 |
| Ilha Grande mangroves | 339 | 3.21E+09 | 0.49 | 0.16 | -9.3 | -0.06 | 0.1 | 58 | 30.99 |
| Interior Alaska/Yukon lowland taiga | 345 | 4.44E+11 | 0.32 | 0.32 | -0.3 | 0 | 0.06 | 11 | 28.67 |
| Interior Yukon/Alaska alpine tundra | 346 | 2.33E+11 | 0.17 | 0.18 | -0.2 | 0 | 0.07 | 13 | 28.42 |
| Iquitos varzea | 347 | 1.15E+11 | 0.2 | 0.18 | -0.7 | -0.12 | 0.27 | 239 | 34.37 |
| Isthmian-Atlantic moist forests | 352 | 5.90E+10 | 0.33 | 0.05 | -7 | -0.48 | 0.28 | 204 | 33.31 |
| Isthmian-Pacific moist forests | 353 | 2.94E+10 | 0.1 | 0.03 | -5.8 | -0.29 | 0.26 | 156 | 33.09 |
| Jalisco dry forests | 356 | 2.62E+10 | 0.09 | 0.01 | -8.4 | -0.54 | 0.11 | 38 | 29.7 |
| Jamaican dry forests | 357 | 2.32E+09 | 0.2 | 0 | -21.8 | -0.37 | 0.02 | 2 | 28.31 |
| Jamaican moist forests | 358 | 8.31E+09 | 0.14 | 0 | -9.9 | 0.36 | 0.03 | 3 | 28.46 |
| Japura-Solimoes-Negro moist forests | 360 | 2.70E+11 | 0.41 | 0.4 | -0.2 | -0.04 | 0.22 | 172 | 33.28 |
| Jurua-Purus moist forests | 366 | 2.43E+11 | 0.17 | 0.17 | -0.1 | -0.01 | 0.23 | 145 | 33.57 |
| Klamath-Siskiyou forests | 387 | 5.04E+10 | 0.15 | 0.09 | -3.5 | 0.01 | 0.06 | 13 | 27.47 |
| La Costa xeric shrublands | 395 | 6.86E+10 | 0.14 | 0 | -13.2 | -1.27 | 0.14 | 90 | 31.23 |
| Lara-Falcon dry forests | 398 | 1.70E+10 | 0.16 | 0 | -10 | -0.78 | 0.09 | 39 | 30.3 |
| Leeward Islands dry forests | 399 | 1.49E+08 | 0.01 | 0 | -4 | 0 | 0.08 | 6 | 27.79 |
| Leeward Islands moist forests | 400 | 9.92E+08 | 0.78 | 0 | -21.6 | -0.8 | 0.04 | 4 | 27.05 |
| Leeward Islands xeric scrub | 401 | 1.65E+09 | 0.25 | 0 | -27.3 | 0.33 | 0.06 | 6 | 27.4 |
| Lesser Antilles mangroves | 402 | 6.53E+08 | 0.31 | 0 | -25.5 | 0.15 | 0.05 | 7 | 27.46 |
| Llanos | 404 | 3.90E+11 | 0.08 | 0.05 | -3.2 | -0.38 | 0.18 | 174 | 32.56 |
| Low Arctic tundra | 407 | 7.98E+11 | 0.17 | 0.17 | 0 | 0 | 0.04 | 6 | 28.35 |
| Madeira-Tapajos moist forests | 422 | 7.21E+11 | 0.27 | 0.27 | -0.3 | -0.02 | 0.2 | 193 | 32.89 |
| Magdalena-Santa Marta mangroves | 425 | 3.20E+09 | 0.27 | 0.01 | -9 | -0.16 | 0.2 | 107 | 31.8 |
| Magdalena-Uraba moist forests | 426 | 7.69E+10 | 0.08 | 0 | -11.7 | -0.54 | 0.22 | 145 | 32.47 |
| Magdalena Valley dry forests | 423 | 1.97E+10 | 0.02 | 0 | -11 | -0.21 | 0.23 | 159 | 32.23 |
| Magdalena Valley montane forests | 424 | 1.05E+11 | 0.15 | 0 | -10 | -0.24 | 0.26 | 281 | 33.52 |
| Magellanic subpolar forests | 427 | 1.48E+11 | 0.54 | 0.52 | -1.3 | -0.1 | 0.09 | 11 | 30.64 |
| Maracaibo dry forests | 437 | 3.03E+10 | 0.15 | 0 | -12.2 | -0.36 | 0.15 | 90 | 31.54 |
| Marajo Varzea forests | 438 | 8.89E+10 | 0.66 | 0.57 | -2.2 | -0.27 | 0.2 | 150 | 32.57 |
| Maranhao Babacu forests | 439 | 1.43E+11 | 0.12 | 0.04 | -6.3 | -0.27 | 0.13 | 67 | 31.35 |
| Maranhao mangroves | 440 | 1.13E+10 | 0.88 | 0.23 | -7.4 | -0.63 | 0.15 | 75 | 31.38 |
| Maranon dry forests | 441 | 1.14E+10 | 0.06 | 0.01 | -4.3 | -0.35 | 0.26 | 192 | 34.05 |
| Marismas Nacionales-San Blas mangroves | 444 | 2.04E+09 | 0.91 | 0.2 | -7 | -0.22 | 0.09 | 17 | 29.07 |
| Mato Grosso seasonal forests | 448 | 4.15E+11 | 0.08 | 0.06 | -1.9 | -0.09 | 0.19 | 141 | 32.7 |
| Mayan Corridor mangroves | 450 | 4.10E+09 | 0.82 | 0.64 | -1.9 | -0.1 | 0.16 | 44 | 31.3 |
| Meseta Central matorral | 458 | 1.26E+11 | 0.05 | 0.02 | -6.1 | -0.11 | 0.12 | 41 | 29.86 |
| Mexican South Pacific Coast mangroves | 460 | 1.17E+09 | 0.08 | 0 | -12.5 | -0.25 | 0.09 | 20 | 29.66 |
| Mid-Continental Canadian forests | 461 | 3.69E+11 | 0.18 | 0.17 | -0.9 | 0.01 | 0.05 | 14 | 28.15 |
| Middle Arctic tundra | 462 | 1.04E+12 | 0.05 | 0.05 | 0 | -0.01 | 0.03 | 3 | 28.78 |
| Middle Atlantic coastal forests | 463 | 1.34E+11 | 0.07 | 0.01 | -9.1 | 0.45 | 0.06 | 11 | 28.12 |
| Midwestern Canadian Shield forests | 465 | 5.47E+11 | 0.14 | 0.13 | -0.4 | 0.01 | 0.05 | 12 | 28.12 |
| Miskito pine forests | 469 | 1.90E+10 | 0.16 | 0.04 | -7.3 | -0.98 | 0.25 | 84 | 32.14 |
| Mississippi lowland forests | 470 | 1.13E+11 | 0.07 | 0.01 | -9.7 | 0.45 | 0.07 | 13 | 28.23 |
| Moist Pacific Coast mangroves | 473 | 1.60E+09 | 0.26 | 0.05 | -6.2 | -1.26 | 0.19 | 69 | 31.16 |
| Mojave desert | 474 | 1.31E+11 | 0.42 | 0.18 | -4.9 | 0.04 | 0.08 | 22 | 27.53 |
| Montana Valley and Foothill grasslands | 476 | 8.19E+10 | 0.05 | 0.01 | -15.1 | 0.02 | 0.08 | 23 | 28.19 |
| Monte Alegre varzea | 478 | 6.70E+10 | 0.16 | 0.16 | -0.5 | -0.1 | 0.22 | 196 | 33.12 |
| Mosquita-Nicaraguan Caribbean Coast mangroves | 479 | 4.45E+09 | 0.43 | 0.03 | -7.1 | -1 | 0.25 | 87 | 32.2 |
| Motagua Valley thornscrub | 480 | 2.34E+09 | 0.03 | 0 | -13.1 | 0 | 0.22 | 86 | 31.56 |
| Muskwa/Slave Lake forests | 484 | 2.63E+11 | 0.14 | 0.14 | -0.1 | 0 | 0.04 | 9 | 28.15 |
| Napo moist forests | 491 | 2.52E+11 | 0.2 | 0.19 | -0.9 | -0.18 | 0.3 | 305 | 34.66 |
| Nebraska Sand Hills mixed grasslands | 494 | 6.13E+10 | 0.02 | 0 | -8 | 0 | 0.08 | 13 | 27.66 |
| Negro-Branco moist forests | 495 | 2.13E+11 | 0.29 | 0.29 | -0.2 | -0.05 | 0.21 | 155 | 33.08 |
| New England/Acadian forests | 502 | 2.38E+11 | 0.06 | 0.02 | -5.3 | 0.26 | 0.06 | 14 | 28.36 |
| Newfoundland Highland forests | 504 | 1.64E+10 | 0.18 | 0.17 | -0.4 | 0 | 0.05 | 8 | 27.81 |
| North Central Rockies forests | 513 | 2.46E+11 | 0.24 | 0.2 | -1.8 | 0.04 | 0.08 | 22 | 28.13 |
| Northeastern Brazil restingas | 523 | 1.01E+10 | 0.3 | 0.12 | -6.3 | -0.56 | 0.08 | 28 | 29.85 |
| Northeastern coastal forests | 527 | 8.98E+10 | 0.04 | 0 | -22.4 | 2.89 | 0.05 | 11 | 28.24 |
| Northern Andean paramo | 530 | 3.00E+10 | 0.42 | 0.06 | -7.3 | -0.21 | 0.29 | 394 | 33.89 |
| Northern California coastal forests | 532 | 1.33E+10 | 0.16 | 0 | -9.2 | 0.42 | 0.05 | 11 | 27.05 |
| Northern Canadian Shield taiga | 533 | 6.15E+11 | 0.07 | 0.07 | 0 | 0 | 0.04 | 7 | 28.15 |
| Northern Cordillera forests | 535 | 2.63E+11 | 0.19 | 0.19 | -0.2 | 0 | 0.05 | 11 | 28.15 |
| Northern Dry Pacific Coast mangroves | 536 | 1.06E+09 | 0.58 | 0 | -16 | 0 | 0.06 | 12 | 28.73 |
| Northern Honduras mangroves | 537 | 1.06E+09 | 0.64 | 0.15 | -7 | -0.73 | 0.26 | 81 | 32.37 |
| Northern mixed grasslands | 549 | 2.19E+11 | 0.08 | 0 | -12.3 | 0.05 | 0.08 | 24 | 28.43 |
| Northern Pacific coastal forests | 542 | 6.06E+10 | 0.4 | 0.34 | -1.1 | -0.07 | 0.04 | 8 | 27.73 |
| Northern short grasslands | 550 | 6.40E+11 | 0.06 | 0.01 | -8.6 | -0.02 | 0.09 | 27 | 28.47 |
| Northern tall grasslands | 551 | 7.62E+10 | 0.08 | 0.02 | -8.6 | 0.35 | 0.06 | 16 | 28.15 |
| Northern transitional alpine forests | 552 | 2.57E+10 | 0.05 | 0.05 | -0.4 | 0.01 | 0.05 | 9 | 27.54 |
| Northwest Mexican Coast mangroves | 555 | 4.99E+09 | 0.47 | 0.17 | -4.8 | -0.06 | 0.08 | 18 | 28.09 |
| Northwest Territories taiga | 557 | 3.47E+11 | 0.08 | 0.08 | -0.1 | -0.04 | 0.05 | 9 | 28.24 |
| Northwestern Andean montane forests | 558 | 8.14E+10 | 0.14 | 0.03 | -5.6 | -0.49 | 0.3 | 324 | 34.06 |
| Oaxacan montane forests | 566 | 7.62E+09 | 0.02 | 0.01 | -6.1 | 0.02 | 0.2 | 91 | 31.36 |
| Ogilvie/MacKenzie alpine tundra | 568 | 2.09E+11 | 0.11 | 0.11 | -0.1 | 0 | 0.05 | 9 | 28.15 |
| Okanogan dry forests | 569 | 5.34E+10 | 0.14 | 0.04 | -8.9 | 0.24 | 0.06 | 16 | 27.97 |
| Orinoco Delta swamp forests | 572 | 2.82E+10 | 0.51 | 0.44 | -2.2 | -1.27 | 0.13 | 76 | 31.14 |
| Orinoco wetlands | 573 | 6.03E+09 | 0.3 | 0.17 | -4.1 | -0.76 | 0.09 | 38 | 30.24 |
| Ozark Mountain forests | 575 | 6.22E+10 | 0.14 | 0.01 | -7.3 | 0.05 | 0.07 | 10 | 27.73 |
| Pacific Coastal Mountain icefields and tundra | 576 | 1.07E+11 | 0.42 | 0.42 | -0.2 | 0 | 0.05 | 9 | 27.76 |
| Palouse grasslands | 579 | 4.70E+10 | 0.03 | 0.01 | -7.4 | 0.41 | 0.08 | 19 | 28.34 |
| Panamanian dry forests | 581 | 5.12E+09 | 0.01 | 0 | -4 | -0.45 | 0.27 | 144 | 33.01 |
| Pantanal | 583 | 1.71E+11 | 0.03 | 0.02 | -1.9 | -0.04 | 0.11 | 64 | 30.98 |
| Pantanos de Centla | 584 | 1.72E+10 | 0.28 | 0.12 | -5.5 | -0.16 | 0.2 | 61 | 31.44 |
| Para mangroves | 585 | 4.42E+09 | 0.48 | 0.11 | -7.9 | -0.99 | 0.16 | 88 | 31.58 |
| Paraguana xeric scrub | 586 | 1.60E+10 | 0.12 | 0 | -16.9 | -0.9 | 0.13 | 65 | 30.95 |
| Parana flooded savanna | 587 | 3.90E+10 | 0.24 | 0.06 | -7.8 | -0.1 | 0.04 | 15 | 29.52 |
| Patagonian grasslands | 589 | 6.33E+10 | 0.04 | 0.03 | -4.8 | -0.26 | 0.09 | 11 | 30.25 |
| Patagonian steppe | 590 | 4.88E+11 | 0.07 | 0.04 | -4.3 | -0.24 | 0.07 | 13 | 30.11 |
| Patia Valley dry forests | 591 | 2.28E+09 | 0 | 0 | -14 | 0 | 0.29 | 146 | 34.28 |
| Pernambuco coastal forests | 595 | 1.76E+10 | 0.04 | 0 | -17.1 | -1.47 | 0.09 | 37 | 30.71 |
| Pernambuco interior forests | 596 | 2.27E+10 | 0.02 | 0 | -16.6 | -1.54 | 0.09 | 38 | 30.71 |
| Peruvian Yungas | 598 | 1.87E+11 | 0.15 | 0.12 | -2.4 | -0.31 | 0.27 | 362 | 34.3 |
| Peten-Veracruz moist forests | 600 | 1.49E+11 | 0.18 | 0.05 | -6.3 | -0.43 | 0.21 | 114 | 31.67 |
| Petenes mangroves | 599 | 1.98E+09 | 0.94 | 0.53 | -4 | -0.1 | 0.07 | 14 | 29.59 |
| Piney Woods forests | 603 | 1.41E+11 | 0.03 | 0 | -8.7 | 0.16 | 0.06 | 11 | 28.15 |
| Puerto Rican dry forests | 607 | 1.28E+09 | 0.08 | 0 | -15 | 3.66 | 0.05 | 6 | 26.76 |
| Puerto Rican moist forests | 608 | 7.55E+09 | 0.05 | 0 | -18.5 | 2.52 | 0.05 | 7 | 26.96 |
| Puget lowland forests | 609 | 2.26E+10 | 0.06 | 0 | -14.9 | 0.73 | 0.05 | 12 | 27.39 |
| Purus-Madeira moist forests | 611 | 1.74E+11 | 0.51 | 0.5 | -0.2 | -0.01 | 0.22 | 162 | 33.4 |
| Purus varzea | 610 | 1.78E+11 | 0.29 | 0.3 | -0.3 | 0 | 0.23 | 190 | 33.45 |
| Queen Charlotte Islands | 618 | 9.98E+09 | 0.47 | 0.42 | -0.9 | -0.29 | 0.04 | 5 | 27.44 |
| Rio Lagartos mangroves | 628 | 3.47E+09 | 0.47 | 0.24 | -5.2 | 0.29 | 0.14 | 32 | 30.7 |
| Rio Negro-Rio San Sun mangroves | 630 | 4.78E+08 | 0.95 | 0.48 | -4.2 | -1.44 | 0.28 | 86 | 33.01 |
| Rio Negro campinarana | 629 | 8.10E+10 | 0.36 | 0.37 | 0 | 0 | 0.22 | 137 | 33.24 |
| Rio Piranhas mangroves | 631 | 2.12E+09 | 0.2 | 0 | -22.2 | -1.7 | 0.07 | 27 | 30.31 |
| Rio Sao Francisco mangroves | 632 | 2.62E+09 | 0.14 | 0.01 | -20.1 | -1.28 | 0.09 | 39 | 30.81 |
| San Lucan xeric scrub | 643 | 3.89E+09 | 0.18 | 0.07 | -8.7 | -0.99 | 0.08 | 9 | 27.49 |
| Santa Marta montane forests | 644 | 4.80E+09 | 0.46 | 0 | -7.4 | -0.64 | 0.15 | 72 | 30.93 |
| Santa Marta paramo | 645 | 1.25E+09 | 0.98 | 0 | -7.1 | -0.53 | 0.15 | 64 | 30.86 |
| Sechura desert | 655 | 1.85E+11 | 0.04 | 0.01 | -6.1 | -0.35 | 0.09 | 31 | 30.26 |
| Semi-arid Pampas | 657 | 3.28E+11 | 0.01 | 0.01 | -6.1 | -0.16 | 0.05 | 13 | 29.2 |
| Serra do Mar coastal forests | 660 | 1.05E+11 | 0.33 | 0.09 | -11.1 | -0.24 | 0.11 | 76 | 31.32 |
| Sierra de la Laguna dry forests | 669 | 3.99E+09 | 0.23 | 0.09 | -7 | -1.02 | 0.08 | 9 | 27.49 |
| Sierra de la Laguna pine-oak forests | 670 | 1.07E+09 | 0.87 | 0.67 | -2.9 | -0.12 | 0.07 | 7 | 27.34 |
| Sierra de los Tuxtlas | 671 | 3.91E+09 | 0.39 | 0 | -8.5 | -0.19 | 0.18 | 52 | 31.36 |
| Sierra Juarez & San Pedro Martir pine-oak forests | 662 | 4.01E+09 | 0.17 | 0.14 | -1.3 | -0.01 | 0.06 | 10 | 27.05 |
| Sierra Madre de Chiapas moist forest | 665 | 1.13E+10 | 0.17 | 0.08 | -4.3 | -0.22 | 0.16 | 55 | 30.24 |
| Sierra Madre de Oaxaca pine-oak forests | 666 | 1.44E+10 | 0.06 | 0.01 | -11.7 | -0.48 | 0.2 | 98 | 31.3 |
| Sierra Madre del Sur pine-oak forests | 667 | 6.13E+10 | 0.03 | 0.01 | -5.6 | -0.31 | 0.13 | 49 | 30.39 |
| Sierra Madre Occidental pine-oak forests | 663 | 2.23E+11 | 0.14 | 0.08 | -3.9 | -0.12 | 0.11 | 44 | 29.4 |
| Sierra Madre Oriental pine-oak forests | 664 | 6.58E+10 | 0.32 | 0.15 | -5.1 | -0.11 | 0.13 | 55 | 29.54 |
| Sierra Nevada forests | 668 | 5.30E+10 | 0.3 | 0.24 | -1.9 | 0.03 | 0.06 | 16 | 27.68 |
| Sinaloan dry forests | 673 | 7.77E+10 | 0.11 | 0.04 | -6.6 | -0.24 | 0.11 | 42 | 29.59 |
| Sinu Valley dry forests | 674 | 2.50E+10 | 0.03 | 0 | -8.9 | -0.52 | 0.17 | 93 | 31.3 |
| Snake/Columbia shrub steppe | 675 | 2.19E+11 | 0.15 | 0.02 | -7.6 | 0.03 | 0.08 | 20 | 28.31 |
| Solimoes-Japura moist forest | 678 | 1.68E+11 | 0.24 | 0.25 | -0.4 | -0.11 | 0.23 | 150 | 33.6 |
| Sonoran-Sinaloan transition subtropical dry forest | 683 | 5.11E+10 | 0.05 | 0.02 | -6.4 | -0.2 | 0.12 | 34 | 29.06 |
| Sonoran desert | 682 | 2.24E+11 | 0.16 | 0.1 | -3.8 | 0 | 0.09 | 29 | 27.96 |
| South Avalon-Burin oceanic barrens | 685 | 2.03E+09 | 0.04 | 0.04 | -1 | -0.04 | 0.06 | 6 | 28.29 |
| South Central Rockies forests | 686 | 1.60E+11 | 0.29 | 0.23 | -1.8 | 0.02 | 0.09 | 24 | 28.4 |
| South Florida rocklands | 690 | 2.08E+09 | 0.11 | 0.01 | -16.6 | 1.37 | 0.04 | 5 | 26.79 |
| Southeastern conifer forests | 707 | 2.37E+11 | 0.05 | 0.01 | -9.9 | 0.36 | 0.05 | 9 | 27.67 |
| Southeastern mixed forests | 708 | 3.49E+11 | 0.02 | 0 | -12.3 | 0.56 | 0.06 | 13 | 28.34 |
| Southern Andean steppe | 714 | 1.79E+11 | 0.17 | 0.11 | -2.8 | -0.16 | 0.08 | 18 | 30.16 |
| Southern Andean Yungas | 713 | 6.13E+10 | 0.06 | 0.02 | -6.1 | -0.2 | 0.1 | 62 | 30.99 |
| Southern Cone Mesopotamian savanna | 716 | 7.78E+10 | 0.02 | 0 | -6.2 | -0.06 | 0.06 | 26 | 30.18 |
| Southern Dry Pacific Coast mangroves | 718 | 9.08E+08 | 0.16 | 0 | -16.3 | -0.38 | 0.17 | 56 | 30.74 |
| Southern Great Lakes forests | 719 | 2.45E+11 | 0.01 | 0 | -15 | 2.09 | 0.04 | 10 | 28.38 |
| Southern Hudson Bay taiga | 720 | 3.75E+11 | 0.18 | 0.19 | -0.1 | 0 | 0.04 | 9 | 28.15 |
| Southern Pacific dry forests | 726 | 4.25E+10 | 0.04 | 0.01 | -9.5 | -0.06 | 0.19 | 95 | 31.23 |
| Southwest Amazon moist forests | 730 | 7.51E+11 | 0.19 | 0.18 | -0.8 | -0.13 | 0.25 | 349 | 34.03 |
| Talamancan montane forests | 763 | 1.64E+10 | 0.57 | 0.27 | -4.7 | -0.25 | 0.27 | 173 | 33.2 |
| Tamaulipan matorral | 764 | 1.63E+10 | 0.06 | 0.03 | -7.1 | -0.73 | 0.09 | 23 | 28.68 |
| Tamaulipan mezquital | 765 | 1.42E+11 | 0.11 | 0.02 | -14.6 | -0.07 | 0.07 | 20 | 28.58 |
| Tapajos-Xingu moist forests | 766 | 3.37E+11 | 0.48 | 0.44 | -0.7 | -0.05 | 0.2 | 147 | 32.86 |
| Tehuacan Valley matorral | 771 | 9.91E+09 | 0.16 | 0.01 | -9.5 | -0.15 | 0.12 | 43 | 29.76 |
| Tehuantepec-El Manchon mangroves | 772 | 2.70E+09 | 0.44 | 0.1 | -7.1 | -0.28 | 0.09 | 19 | 29.11 |
| Tepuis | 774 | 4.90E+10 | 0.63 | 0.64 | -0.2 | -0.1 | 0.2 | 156 | 32.58 |
| Texas blackland prairies | 776 | 5.04E+10 | 0.01 | 0 | -13.7 | 0 | 0.06 | 11 | 27.73 |
| Tocantins/Pindare moist forests | 786 | 1.94E+11 | 0.11 | 0.04 | -5.1 | -0.46 | 0.16 | 97 | 31.79 |
| Torngat Mountain tundra | 790 | 3.24E+10 | 0.42 | 0.43 | 0 | 0 | 0.05 | 4 | 28.12 |
| Trans-Mexican Volcanic Belt pine-oak forests | 794 | 9.20E+10 | 0.18 | 0.02 | -12.7 | -0.16 | 0.14 | 66 | 30.47 |
| Trinidad and Tobago dry forests | 796 | 2.72E+08 | 0.05 | 0 | -20.1 | -2.46 | 0.06 | 15 | 29.17 |
| Trinidad and Tobago moist forests | 797 | 4.75E+09 | 0.3 | 0 | -15.4 | -3.23 | 0.06 | 16 | 29.25 |
| Trinidad mangroves | 798 | 1.86E+08 | 0.15 | 0 | -25.2 | -4.45 | 0.06 | 15 | 29.22 |
| Tumbes-Piura dry forests | 803 | 4.14E+10 | 0.07 | 0.02 | -5.1 | -0.16 | 0.21 | 123 | 32.13 |
| Uatuma-Trombetas moist forests | 805 | 4.74E+11 | 0.45 | 0.44 | -0.2 | -0.02 | 0.22 | 190 | 33.04 |
| Ucayali moist forests | 806 | 1.15E+11 | 0.21 | 0.2 | -1.5 | -0.47 | 0.3 | 331 | 34.78 |
| Upper Midwest Forest/Savanna transition zone | 808 | 1.66E+11 | 0.04 | 0 | -14.4 | 1.46 | 0.08 | 19 | 28.54 |
| Uruguayan savanna | 810 | 3.56E+11 | 0.03 | 0.01 | -7.5 | -0.07 | 0.07 | 37 | 30.66 |
| Usumacinta mangroves | 812 | 3.14E+09 | 0.83 | 0.47 | -5.2 | -0.27 | 0.15 | 38 | 30.94 |
| Valdivian temperate forests | 813 | 2.49E+11 | 0.23 | 0.16 | -3.1 | -0.3 | 0.08 | 14 | 30.39 |
| Venezuelan Andes montane forests | 815 | 2.95E+10 | 0.5 | 0 | -11.6 | -0.23 | 0.18 | 132 | 31.97 |
| Veracruz dry forests | 816 | 6.65E+09 | 0.05 | 0 | -14.4 | -0.42 | 0.17 | 49 | 31.17 |
| Veracruz moist forests | 817 | 6.93E+10 | 0.08 | 0.02 | -6.9 | -0.08 | 0.13 | 52 | 29.98 |
| Veracruz montane forests | 818 | 4.97E+09 | 0.06 | 0 | -20.3 | 0.75 | 0.13 | 43 | 30.03 |
| Wasatch and Uinta montane forests | 823 | 4.16E+10 | 0.1 | 0.06 | -4.6 | 0.28 | 0.06 | 15 | 27.97 |
| Western Ecuador moist forests | 830 | 3.42E+10 | 0.04 | 0 | -7.5 | -0.39 | 0.28 | 184 | 33.85 |
| Western Great Lakes forests | 832 | 2.75E+11 | 0.16 | 0.07 | -5.2 | 0.21 | 0.06 | 15 | 28.31 |
| Western Gulf coastal grasslands | 834 | 8.08E+10 | 0.12 | 0.03 | -8.8 | 0.21 | 0.05 | 12 | 28.26 |
| Western short grasslands | 843 | 4.36E+11 | 0.01 | 0 | -9 | 0.04 | 0.09 | 28 | 28.34 |
| Willamette Valley forests | 845 | 1.49E+10 | 0.02 | 0 | -19.6 | 1.8 | 0.04 | 8 | 27.39 |
| Windward Islands dry forests | 846 | 4.93E+08 | 0.15 | 0 | -27.4 | -1.7 | 0.05 | 5 | 27.73 |
| Windward Islands moist forests | 847 | 2.02E+09 | 0.39 | 0.01 | -17.4 | -0.85 | 0.08 | 9 | 28.08 |
| Windward Islands xeric scrub | 848 | 1.03E+09 | 0.2 | 0 | -29.9 | -1.58 | 0.07 | 8 | 27.73 |
| Wyoming Basin shrub steppe | 850 | 1.33E+11 | 0.04 | 0 | -8.4 | -0.01 | 0.07 | 17 | 28.11 |
| Xingu-Tocantins-Araguaia moist forests | 851 | 2.67E+11 | 0.15 | 0.14 | -0.7 | -0.09 | 0.18 | 126 | 32.25 |
| Yucatan dry forests | 856 | 4.99E+10 | 0.1 | 0.03 | -7.8 | 0.02 | 0.13 | 33 | 30.7 |
| Yucatan moist forests | 857 | 6.99E+10 | 0.23 | 0.19 | -2.1 | -0.11 | 0.19 | 61 | 31.42 |
| Yukon Interior dry forests | 858 | 6.25E+10 | 0.02 | 0.02 | -0.8 | 0 | 0.05 | 9 | 27.81 |
| Zacatonal | 860 | 3.03E+08 | 0.61 | 0.28 | -5.2 | -0.38 | 0.12 | 47 | 29.76 |

Table S3: List of the 139 threatened high-sensitivity species with no or minor coverage by intact protected habitat. Intact protected habitat corresponds to the area within each species’ range that is both protected and with a human footprint <4 (see Methods). The proportion of intact protected habitat is obtained by dividing by the species’ range size.

| **Name** | **Sensitivity** | **Area of intact protected habitat**  **(km^2^)** | **Proportion of intact protected habitat (%)** | **RL**  **status** |
| --- | --- | --- | --- | --- |
| *Acrobatornis fonsecai* | 43.8 | 32 | 0.86 | VU |
| *Agelaius xanthomus* | 43.6 | 0 | 0 | EN |
| *Amazilia alfaroana* | 46 | 0 | 0 | CR |
| *Amazona imperialis* | 38.6 | 0 | 0 | CR |
| *Ampelornis griseiceps* | 40 | 489 | 2.72 | VU |
| *Anairetes alpinus* | 39 | 611 | 3.05 | EN |
| *Antilophia bokermanni* | 46.4 | 3 | 9.16 | CR |
| *Antrostomus noctitherus* | 43.3 | 0 | 0 | EN |
| *Anumara forbesi* | 39.4 | 0 | 0 | EN |
| *Asthenes perijana* | 39.5 | 0 | 0 | EN |
| *Atlapetes blancae* | 40.4 | 0 | 0 | CR |
| *Automolus lammi* | 46.9 | 87 | 0.14 | EN |
| *Bangsia aureocincta* | 44.9 | 25 | 2.07 | EN |
| *Bangsia melanochlamys* | 43 | 14 | 0.35 | VU |
| *Basileuterus griseiceps* | 45.3 | 7 | 0.77 | EN |
| *Buteo ridgwayi* | 39.5 | 0 | 0 | CR |
| *Campephilus principalis* | 44.6 | 0 | 0 | CR |
| *Capito hypoleucus* | 42.4 | 0 | 0 | VU |
| *Capito quinticolor* | 42.1 | 598 | 1.47 | VU |
| *Catharopeza bishopi* | 39.6 | 0 | 0 | EN |
| *Celeus tinnunculus* | 44.2 | 1592 | 1.32 | VU |
| *Centrocercus minimus* | 46.2 | 249 | 2.57 | EN |
| *Chasiempis ibidis* | 46 | 2 | 0.85 | EN |
| *Cichlopsis leucogenys* | 38.9 | 1507 | 1.47 | EN |
| *Cinclodes palliatus* | 41.4 | 310 | 1.28 | CR |
| *Cistothorus apolinari* | 40.5 | 7 | 0.26 | EN |
| *Clytoctantes alixii* | 37.8 | 124 | 0.16 | EN |
| *Coccyzus rufigularis* | 37.9 | 0 | 0 | EN |
| *Coeligena orina* | 41.9 | 162 | 8.94 | CR |
| *Corvus leucognaphalus* | 41.7 | 329 | 3.95 | VU |
| *Cotinga maculata* | 45.8 | 816 | 2.68 | CR |
| *Cranioleuca berlepschi* | 39.1 | 172 | 6.61 | VU |
| *Crax alberti* | 50 | 0 | 0 | CR |
| *Crax blumenbachii* | 44.5 | 110 | 9.52 | EN |
| *Cyanolyca mirabilis* | 45.1 | 1 | 0.01 | VU |
| *Dendrortyx barbatus* | 44.6 | 11 | 0.13 | VU |
| *Diglossa venezuelensis* | 44.1 | 7 | 0.87 | EN |
| *Doliornis sclateri* | 39.8 | 363 | 2.77 | VU |
| *Dubusia carrikeri* | 40.3 | 0 | 0 | EN |
| *Eleoscytalopus psychopompus* | 45.4 | 273 | 5.3 | EN |
| *Eriocnemis isabellae* | 40.4 | 0 | 0 | CR |
| *Eriocnemis mirabilis* | 41.5 | 0 | 0 | EN |
| *Eriocnemis nigrivestis* | 38.9 | 1 | 1.27 | CR |
| *Eupherusa cyanophrys* | 43.4 | 1 | 0.02 | EN |
| *Eupherusa poliocerca* | 41.6 | 8 | 0.05 | VU |
| *Formicivora paludicola* | 43.7 | 0 | 0 | CR |
| *Geospiza pauper* | 39.3 | 0 | 0 | CR |
| *Geothlypis speciosa* | 42.5 | 7 | 1.4 | EN |
| *Glaucidium nubicola* | 42.2 | 439 | 6.31 | VU |
| *Glaucis dohrnii* | 40.7 | 71 | 7.22 | EN |
| *Grallaria chthonia* | 45.6 | 0 | 0 | CR |
| *Grallaria excelsa* | 43.6 | 74 | 0.87 | VU |
| *Grallaria fenwickorum* | 48.7 | 0 | 0 | CR |
| *Grallaria kaestneri* | 41.8 | 0 | 0 | EN |
| *Grallaria saltuensis* | 42.9 | 0 | 0 | EN |
| *Grallaricula cucullata* | 45.3 | 331 | 2.13 | VU |
| *Grallaricula cumanensis* | 44.3 | 0 | 0 | VU |
| *Hapalopsittaca fuertesi* | 42.7 | 10 | 0.7 | CR |
| *Hemignathus wilsoni* | 47 | 11 | 6.17 | EN |
| *Hemitriccus kaempferi* | 43.9 | 325 | 4.12 | VU |
| *Hemitriccus mirandae* | 37.7 | 123 | 0.16 | VU |
| *Henicorhina negreti* | 44.2 | 132 | 6.82 | VU |
| *Herpsilochmus parkeri* | 42.4 | 0 | 0 | EN |
| *Hylonympha macrocerca* | 41.3 | 0 | 0 | EN |
| *Hylopezus auricularis* | 45.7 | 0 | 0 | VU |
| *Hylorchilus navai* | 44.7 | 138 | 3.13 | VU |
| *Leistes defilippii* | 38.4 | 0 | 0 | VU |
| *Leptotila conoveri* | 40.1 | 0 | 0 | EN |
| *Leptotila wellsi* | 38.2 | 0 | 0 | CR |
| *Leucopeza semperi* | 43.3 | 0 | 0 | CR |
| *Lipaugus weberi* | 50 | 0 | 0 | CR |
| *Lophornis brachylophus* | 38.3 | 0 | 0 | CR |
| *Loxia megaplaga* | 43.6 | 212 | 5.95 | EN |
| *Loxioides bailleui* | 39.2 | 0 | 0 | CR |
| *Loxops caeruleirostris* | 48.2 | 0 | 0 | CR |
| *Macroagelaius subalaris* | 43.1 | 556 | 1.7 | EN |
| *Megascops barbarus* | 45.3 | 0 | 0 | VU |
| *Megascops gilesi* | 41.8 | 0 | 0 | VU |
| *Merulaxis stresemanni* | 48.9 | 0 | 0 | CR |
| *Myioborus pariae* | 42.2 | 0 | 0 | EN |
| *Myiotheretes pernix* | 40.5 | 0 | 0 | EN |
| *Myrmoderus ruficauda* | 41.2 | 1349 | 1.6 | EN |
| *Myrmotherula urosticta* | 43.3 | 1701 | 1.63 | VU |
| *Nemosia rourei* | 44.7 | 0 | 0 | CR |
| *Nesopsar nigerrimus* | 46.7 | 0 | 0 | EN |
| *Nothoprocta taczanowskii* | 40.2 | 3 | 0.01 | VU |
| *Nothura minor* | 37.8 | 2011 | 0.42 | VU |
| *Odontophorus atrifrons* | 46.6 | 0 | 0 | VU |
| *Odontophorus strophium* | 48 | 25 | 0.72 | VU |
| *Onychorhynchus occidentalis* | 43.7 | 278 | 2.63 | VU |
| *Oreophasis derbianus* | 50 | 655 | 2.1 | EN |
| *Oxypogon cyanolaemus* | 39.9 | 0 | 0 | CR |
| *Pauxi pauxi* | 50 | 410 | 0.37 | EN |
| *Pauxi unicornis* | 48.7 | 0 | 0 | CR |
| *Percnostola arenarum* | 44.2 | 400 | 4.51 | VU |
| *Phibalura boliviana* | 39.1 | 0 | 0 | EN |
| *Phyllomyias urichi* | 44.9 | 17 | 1.57 | EN |
| *Phyllomyias weedeni* | 40.2 | 617 | 4.37 | VU |
| *Phylloscartes beckeri* | 44.7 | 70 | 3.02 | EN |
| *Phylloscartes ceciliae* | 41.8 | 3 | 0.02 | CR |
| *Pipile pipile* | 47 | 0 | 0 | CR |
| *Pipilo socorroensis* | 38.1 | 0 | 0 | EN |
| *Podiceps gallardoi* | 38.8 | 1524 | 3.34 | CR |
| *Pogonotriccus lanyoni* | 40.8 | 0 | 0 | EN |
| *Poospiza goeringi* | 40.9 | 13 | 0.34 | VU |
| *Premnoplex pariae* | 47 | 0 | 0 | EN |
| *Premnoplex tatei* | 42.1 | 0 | 0 | EN |
| *Psarocolius cassini* | 40.2 | 337 | 3.59 | VU |
| *Pseudastur occidentalis* | 39.1 | 190 | 0.38 | EN |
| *Pyrrhura orcesi* | 40.8 | 0 | 0 | EN |
| *Rhynchopsitta terrisi* | 38.8 | 88 | 6.86 | EN |
| *Rollandia microptera* | 39.6 | 35 | 0.09 | EN |
| *Scytalopus canus* | 42.3 | 0 | 0 | EN |
| *Scytalopus gonzagai* | 39.8 | 26 | 0.59 | EN |
| *Scytalopus iraiensis* | 40 | 74 | 1.43 | EN |
| *Scytalopus novacapitalis* | 38.1 | 635 | 0.53 | EN |
| *Scytalopus perijanus* | 40.9 | 0 | 0 | VU |
| *Scytalopus robbinsi* | 45.9 | 0 | 0 | EN |
| *Scytalopus rodriguezi* | 45.9 | 0 | 0 | EN |
| *Setophaga angelae* | 45.5 | 0 | 0 | EN |
| *Setophaga chrysoparia* | 40.4 | 8 | 0.02 | EN |
| *Sitta insularis* | 47.3 | 0 | 0 | CR |
| *Synallaxis courseni* | 39.4 | 0 | 0 | VU |
| *Synallaxis fuscorufa* | 40.4 | 0 | 0 | VU |
| *Synallaxis kollari* | 41.1 | 246 | 5.96 | CR |
| *Synallaxis maranonica* | 39 | 0 | 0 | CR |
| *Tachycineta euchrysea* | 40.5 | 276 | 2.75 | VU |
| *Tangara fastuosa* | 42 | 3 | 0.02 | VU |
| *Terenura sicki* | 43.1 | 0 | 0 | CR |
| *Thryophilus nicefori* | 42.1 | 1 | 0.02 | CR |
| *Thryophilus sernai* | 38.8 | 0 | 0 | EN |
| *Toxostoma guttatum* | 39.8 | 0 | 0 | CR |
| *Troglodytes monticola* | 41.1 | 0 | 0 | CR |
| *Troglodytes tanneri* | 41.8 | 0 | 0 | VU |
| *Tympanuchus cupido* | 48.3 | 110 | 0.03 | VU |
| *Tympanuchus pallidicinctus* | 41 | 98 | 0.15 | VU |
| *Xenoglaux loweryi* | 46.9 | 108 | 6.61 | EN |
| *Xenospiza baileyi* | 38.9 | 0 | 0 | EN |
| *Zaratornis stresemanni* | 42.5 | 726 | 0.91 | VU |
